# A multi-pronged human microglia response to Alzheimer’s disease Aβ pathology

**DOI:** 10.1101/2022.07.07.499139

**Authors:** Renzo Mancuso, Nicola Fattorelli, Anna Martinez-Muriana, Emma Davis, Leen Wolfs, Johanna Van Den Daele, Ivana Geric, Pranav Preman, Lutgarde Serneels, Suresh Poovathingal, Sriram Balusu, Catherine Verfaille, Mark Fiers, Bart De Strooper

**Author notes:** These authors contributed equally: Renzo Mancuso, Nicola Fattorelli, Anna Martinez-Muriana.

## Abstract

Microglial activation and neuroinflammation are initial steps in the pathogenesis of Alzheimer’s disease (AD). However, studies in mouse models and human postmortem samples have yielded divergent results regarding microglia cell states relevant to AD. Here, we investigate 127,000 single cell expression profiles of human microglia isolated freshly from a xenotransplantation model for early AD. While human microglia adopt a disease-associated (DAM) profile, they display a much more pronounced HLA-cell state related to antigen presentation in response to amyloid plaques. In parallel, a distinctive pro-inflammatory cytokine and chemokine CRM response is mounted against oligomeric amyloid-β. TREM2 and, to a lesser extent, APOE polymorphisms, modulate the response of microglia to amyloid-β plaques, in contrast with the response to oligomeric Aβ. Specific polygenic risk genes are enriched in each branch of these multi-pronged response of human microglia to amyloid pathology (ARM). ARM responses can be captured in post-mortem studies when reanalyzed in light of this novel, comprehensive data set. In conclusion, therapeutic strategies targeting microglia in AD need to carefully assess how they affect the different cell states, as the overall balance between distinct microglial profiles might determine a protective or damaging outcome.

---

Microglia are a central part of the cellular response in AD pathogenesis^1,2^, in particular in the early response to amyloid-β pathology^3–6^. Murine microglia transit from a homeostatic state towards “reactive” disease-associated microglia (DAM), also called activated response microglia or neurodegenerative microglia (MGnD)^4,5,7^. This response is partially TREM2-dependent and elicited by fibrillary amyloid-β structures in mice^3,4,8^. Ageing, tauopathy, amyotrophic lateral sclerosis^4,5,9,10^, apoptotic neurons^5^, Danish amyloid^11^ and even peripheral lipid dyshomeostasis^12^ can elicit a similar response, suggesting that the DAM state is a generic response of mouse microglia to damage. However, it is highly controversial whether a DAM response exists in the human brain^13,14^. Mouse and human microglia are evolutionary divergent, especially regarding the expression of relevant AD genetic risk^8,15,16^. In addition, the microglial cell states reported by post-mortem samples are highly inconsistent. Mathys et al.^17^ found 22 genes upregulated in microglia from AD patients (only 5 overlapping with the DAM signature). Grubman et al.^18^ reported 8 genes, whereas Zhou et al.^19^ identified 11 DAM genes enriched in AD patients compared with controls. DelAguila et al.^20^ analysed single-nucleus transcriptomes from 3 AD patients and were unable to recapitulate an activation signature. A large study^21^ analyzed 482,472 single nuclei from non-demented control and AD brains and indicated several distinct transcriptomic states of microglia encompassing DAM-like (*ITGAX, SPP1*) and proinflammatory (*IL1B, NFKB1*) states in AD. The lack of congruency in these human postmortem brain studies can be partially explained by technical issues, such as post-mortem time^13^ or intrinsic lesser resolution of single nuclei sequencing approaches^14^, heterogeneity in pathological samples, or technical shortcomings including sample size. Additionally, there are inherent limitations to post-mortem studies as they only reflect terminal stages of a complex, decade-long disease process, that often encompasses additional concomitant neuro pathologies (TDP-43, vascular dementia, or alpha-synuclein).

Here, we circumvent these multiple drawbacks by using a human microglia xenotransplantation model^22,23^. We provide a full map of the different cell states adopted by human microglia in response to amyloid-β pathology, including the elusive component of soluble oligomeric amyloid-β that have been linked to neuroinflammation, neurodegeneration and cognitive impairment^24–27^. We show that human microglia react very differently from mouse microglia to amyloid-β pathology. This multi-pronged amyloid-β response (ARM) encompasses several branches, all differentially enriched in polygenic risk, and that can be selectively disturbed by introducing *TREM2* or *APOE* polymorphisms.

## A multi-pronged reaction of human microglia to amyloid-β pathology

We transplanted microglia precursors in the mouse brain according to the MIGRATE protocol^23^. We used three different host genetic backgrounds i.e. *Rag2*^*-/-*^ *Il2rγ*^*-/-*^ *hCSF1*^*KI*^ *App*^*NL-G-F*^ (with progressive amyloid-β plaques accumulation from 3 months after birth, thereafter called *App*^*NL-G-F*^), *Rag2*^*-/-*^ *Il2rγ*^*-/-*^ *hCSF1*^*KI*^ *App*^*hu/hu*^ mice (humanized *App* control mice^28^, *App*^*hu/hu*^) and *Rag2*^*-/-*^ *Il2rγ*^*-/-*^ *hCSF1*^*KI*^ *App*^*NL-G-F*^ *ApoE*^*-/-*^ (*App*^*NL-G-F*^ *ApoE*^*-/-*^) mice. We also studied stem cell-derived microglia from three different human genetic backgrounds i.e. UKBIO11-A, BIONi010-C and H9 (see methods Table 1). Microglia were transplanted in 4 days old neonates (P4) and isolated at 6-7 months after transplantation by FACS (CD11b+ hCD45+, Fig. 1a and Extended Data Fig. 1). We obtained high-quality single cell transcriptomes from more than 127,000 microglia across 95 mice, excluding CNS-associated macrophages (CAM), proliferative cells, other myeloid cells and low-quality cells and doublets (Extended Data Fig. 2 and 3). We identified five distinct microglia clusters that we annotate as Homeostatic (HM), Cytokine response-1 and 2 (CRM-1 and -2), Interferon response (IRM), Disease-Associated response (DAM) and Antigen-presenting response (HLA), as well as two transitory clusters that we named Ribosomal microglia (RM) enriched in ribosomal genes, and Transitioning CRM (tCRM) microglia which show high levels of homeostatic genes but also express CRM markers (Fig. 1b-d and Extended Data Fig. 4). The DAM and IRM clusters have similar counterparts in mouse^3,4^ (Extended Data Fig. 4b). CRM and HLA represent novel clusters not identified before in the mouse microglia of transgenic models of amyloid-β pathology. They are defined by the upregulation of genes encoding for cytokines and chemokines (*CCL4, CCL3, CCL42L, CCL3L1, GADD45B, CCL2, CH25H, IL1B, ZFP36, SGK1*), and different elements of the HLA family (*HLA-DRA, HLA-DPA1, HLA-DRB5, CD74, HLA-DPB1, HLA-DMA, MS4A6A, HLA-DMB*), respectively (Fig. 1d and Extended Data Fig. 4). Interestingly, whereas the DAM profile shows a clear downregulation of homeostatic microglia genes, these partially regain expression in CRM, HLA and IRM clusters (Fig. 1c)

**Table 1.**
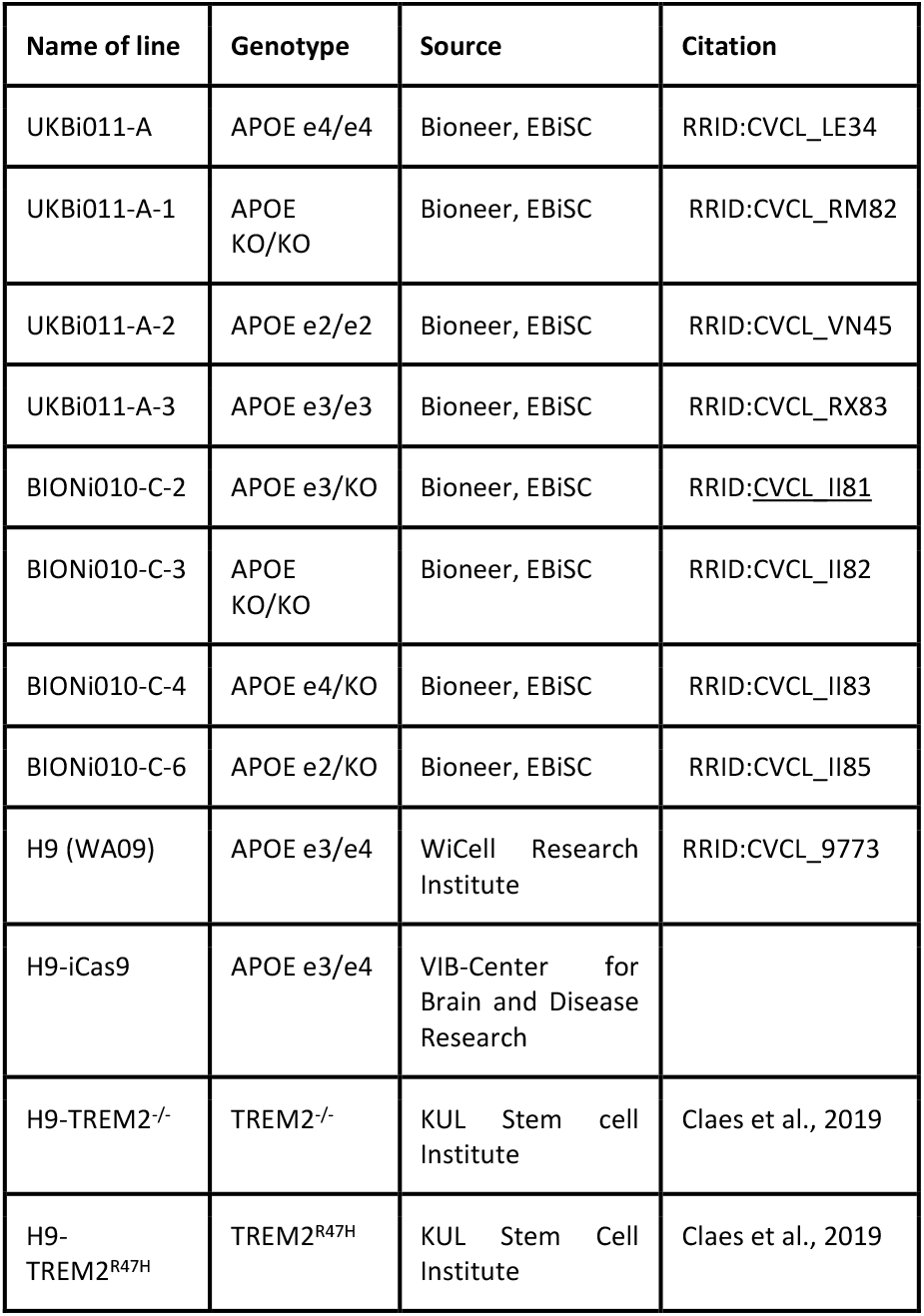
Summary of cell lines used in this study.

**Figure 1.**
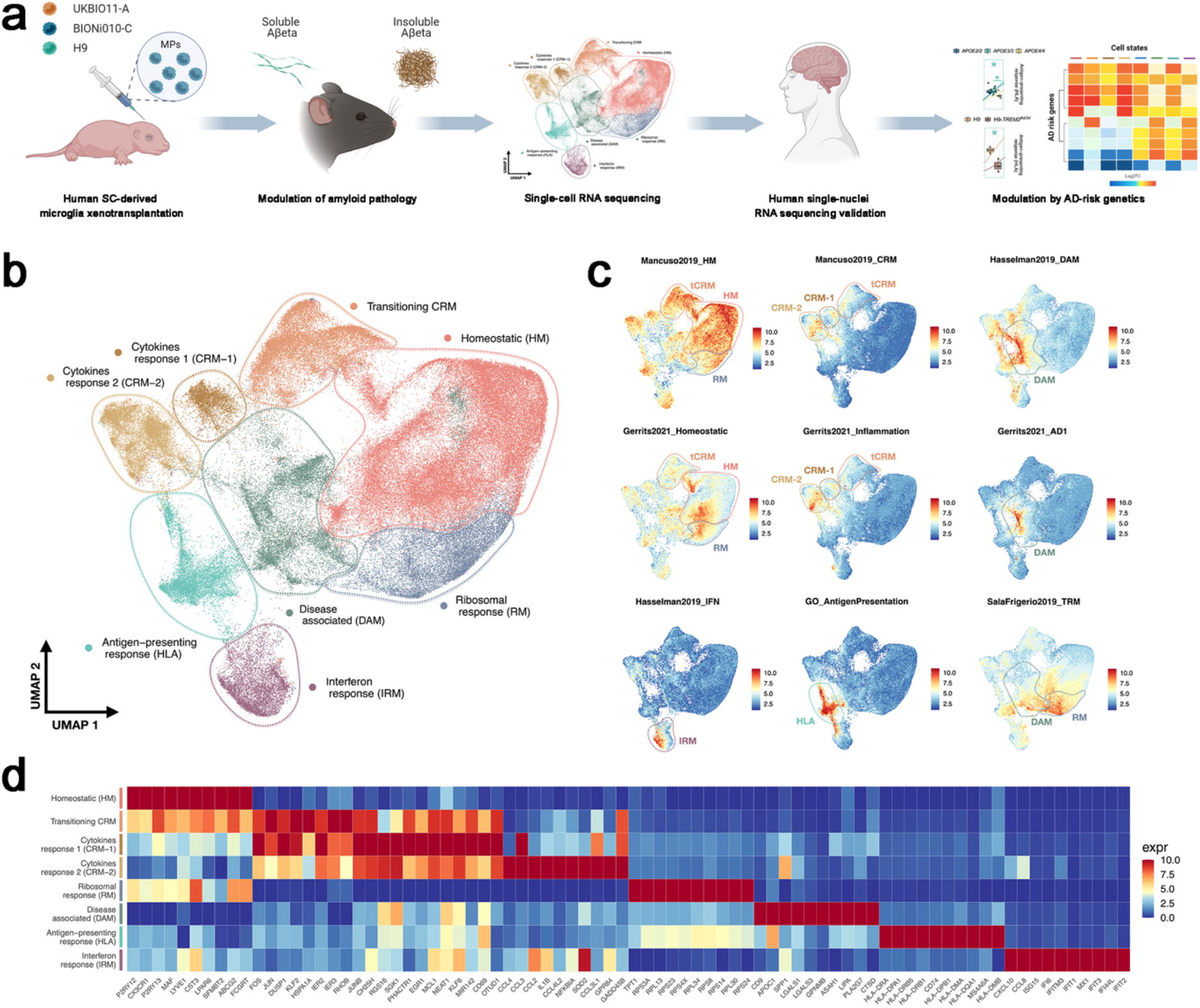
Human microglia display a complex, heterogeneous response to amyloid-β pathology. **a**, Experimental design used in this study. **b**, UMAP plot visualizing 127,755 single xenografted human microglial cells sorted from mouse brain (CD11b^+^ hCD45^+^) ∼6 months after transplantation. Cells are coloured according to clusters identified: homeostatic (HM), Ribosomal response (RM), Disease Associated (DAM), Interferon response (IRM), Antigen presenting response (HLA), Cytokine response 1/2 (CRM-1 and CRM-2), and Transitioning CRM (tCRM). The assignment of different clusters to distinct cell types or states is based on previous experimental data from our and other laboratories^4,7,8,15^ (see c, d and Extended Data Fig. 3 and 4). **c**, UMAP plots as in (b), coloured by the combined level of expression of groups of genes that characterise distinct microglial transcriptional states. **d**, Top 10 most differentially expressed genes in each cluster (normalized expression scaled by gene is shown).

**Figure 2.**
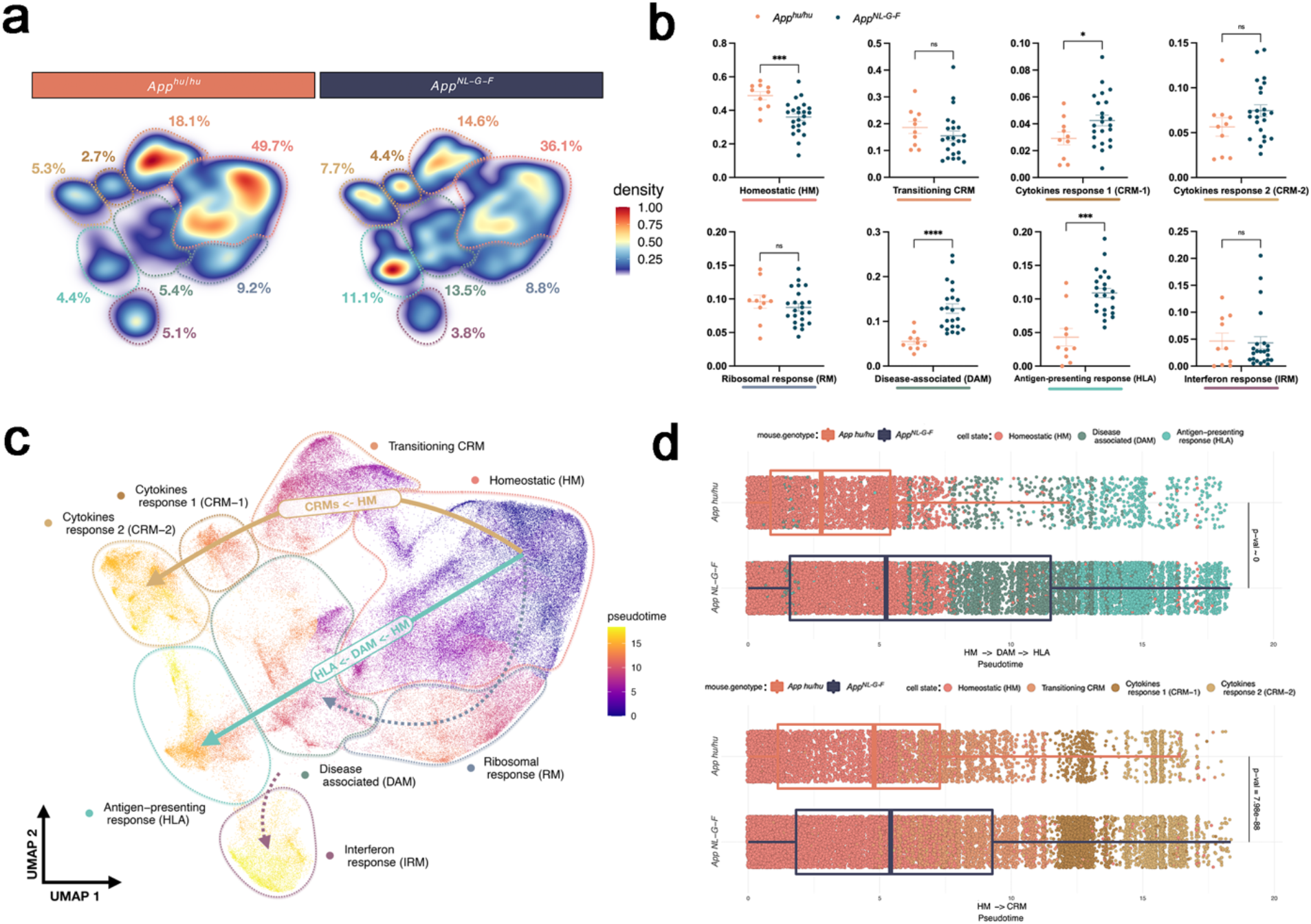
Multi-pronged response of human microglia upon exposure to amyloid-β. **a**, Density plots displaying the average distribution of human microglia transplanted in *App*^*NL-G-F*^ (n=23) and *App*^*hu/hu*^ (n=10) mice (density is normalized to sample size). **b**, Distribution and proportion of cells across all identified clusters. Microglia transplanted in *App*^*NL-G-F*^ mice acquire three distinctive phenotypic states: DAM, HLA and CRM (each dot represents a single mouse, unpaired t-test, *p<0.05, ***p<0.001, ****p<0,0001). **c**, Phenotypic trajectory followed by the human microglia after exposure to amyloid-β *in vivo*, obtained by an unbiased pseudotime ordering with Monocle 3. Note the two distinct transcriptional routes towards either CRM or to DAM and HLA. **d**, Distribution of cell from different host mouse genetic backgrounds (y-axis) across the two main transcriptional trajectories, DAM and HLA (top panel) and CRM (bottom panel), colored by clusters shown in a. Note the shift in transcriptional states in *App*^*NL-G-F*^ vs *App*^*hu/hu*^ mice.

**Figure 3.**
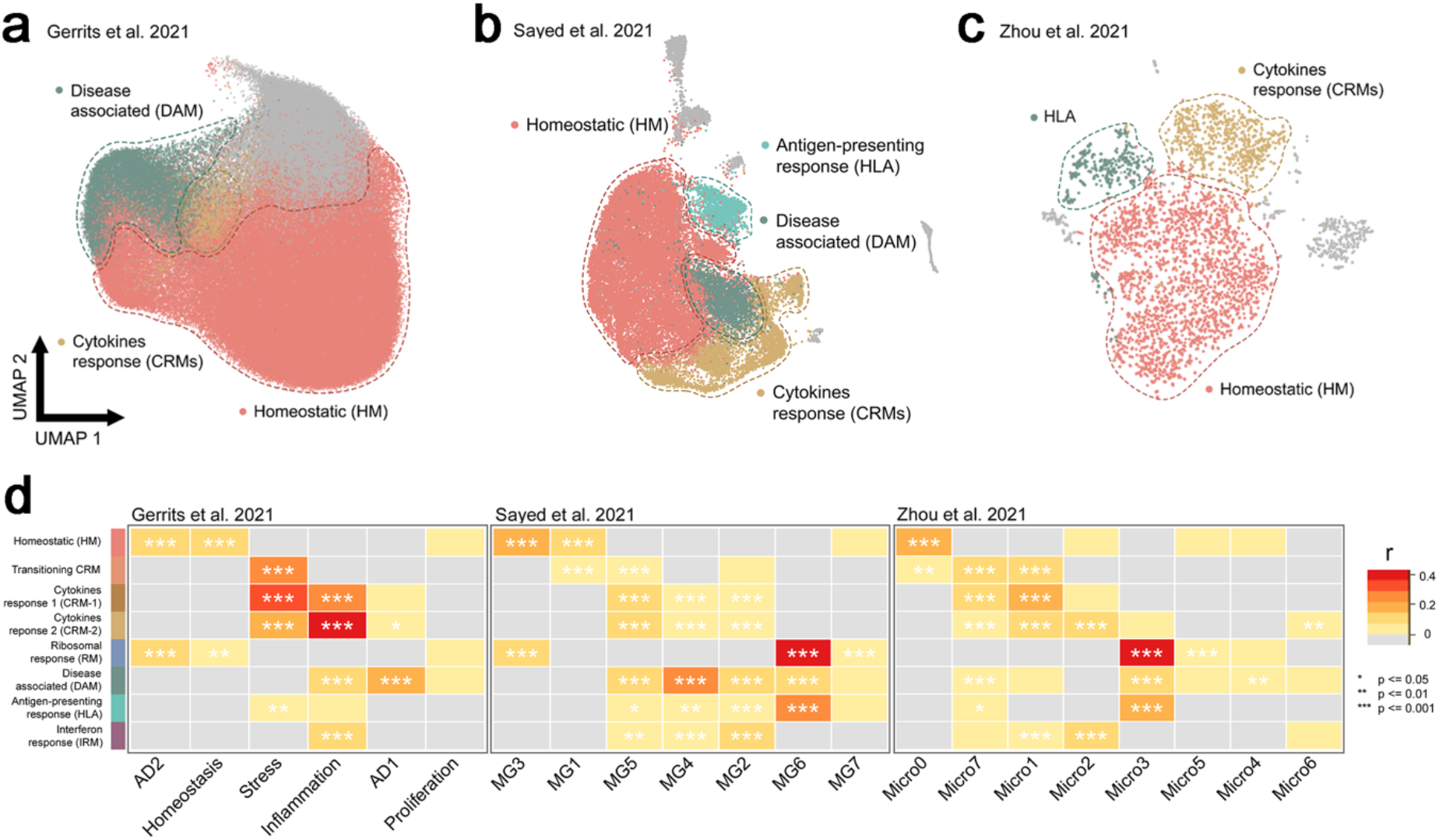
Single microglia nuclei from human postmortem brain. **a**, Seurat ModuleScore calculated for human snRNA-seq studies and annotated using top 100 DE genes (p<0.05) from our xenotransplanted microglia profiles, with resulting module scores displayed on UMAP of reprocessed (see Methods) single nuclei from **a**, Gerrits et al. (n=176,136) **b**, Sayed et al. (n=28,767), and **c**, Zhou et al. (n=3,978). **d**, Pairwise Pearson correlation between logFC of all DE genes (p<0.05) of each microglia sub-type and logFC of all DE genes (p<0.05) of clusters from each human snRNA-seq study, with significance denoted by asterisks. Only positive correlations are depicted here (r>0). Additional correlations are shown in Extended Data Fig. 6 and 7.

**Figure 4.**
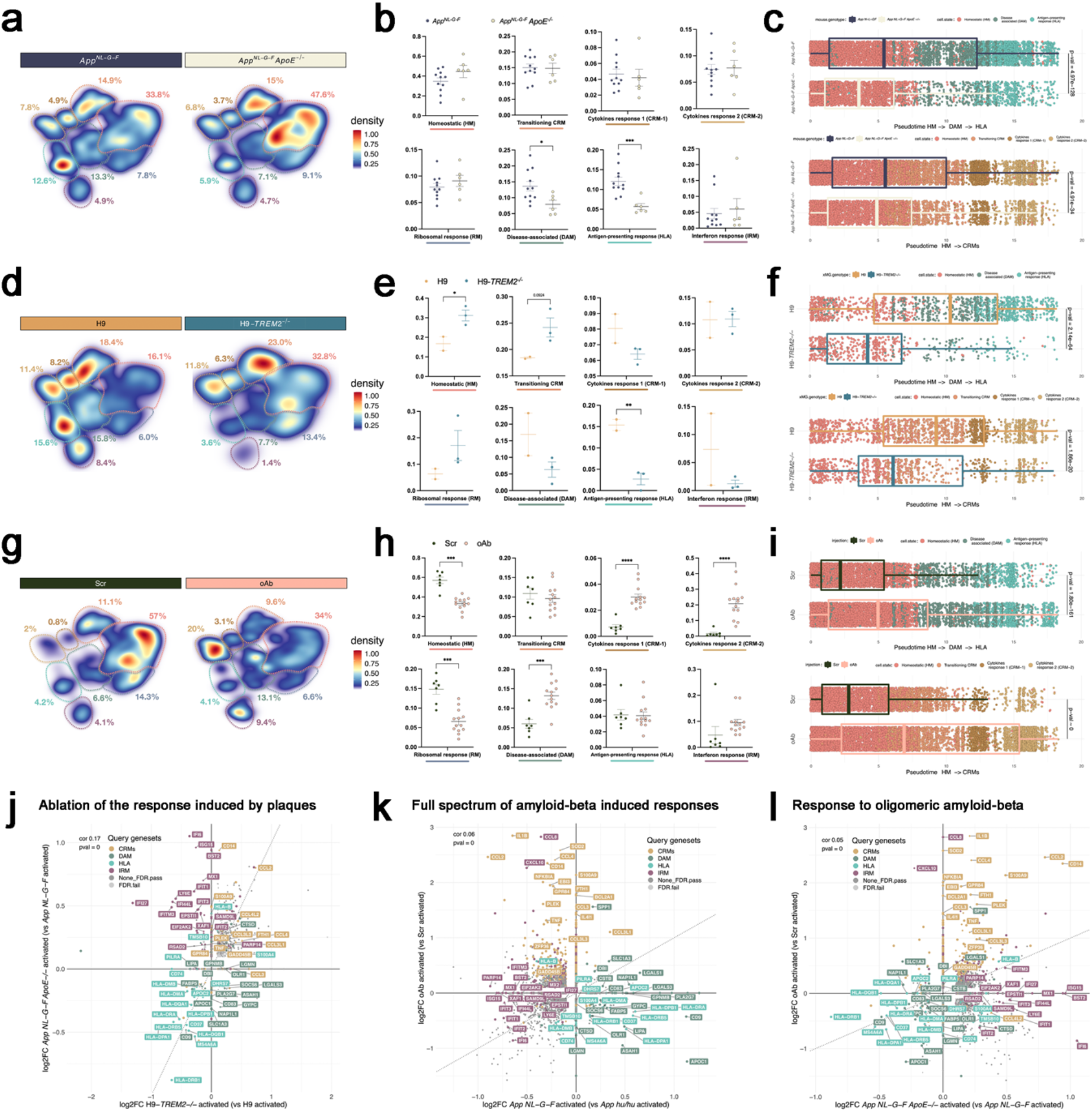
Human microglia display a differential response to amyloid-β plaques and soluble oligomers. **a**, Density plots displaying the average distribution of human microglia transplanted in *App*^*NL-G-F*^ (n=11) and *App*^*NL-G-F*^ *ApoE*^*-/-*^ (n=6) mice (density is normalized by sample size). **b**, Distribution and proportion of cells across all identified clusters. Absence of amyloid-β plaques and ApoE led to a significant reduction in DAM and HLA transcriptional responses (each dot represents a single mouse, unpaired t-test, *p<0.05, ***p<0.01) while CRM responses remain unaffected. **c**, Phenotypic trajectory followed by the human microglia obtained by an unbiased pseudotime ordering with Monocle 3. Proportion of cells from different host mouse genetic backgrounds (y-axis) present at different stages of the pseudotime trajectory (x-axis), colored by clusters shown in Fig. 1a. Note the shift in transcriptional states in *App*^*NL-G-F*^ x *ApoE*^*-/-*^ vs. *App*^*NL-G-F*^ mice consistent with a reduction in both DAM and HLA transcriptional states. **d**, Density plots displaying the average distribution of human H9-*WT* (n=2) and H9-*TREM2*^*-/-*^ (n=3) microglia transplanted in *App*^*NL-G-F*^ mice (density is normalized by sample size). **e**, Distribution and proportion of cells across all identified clusters. Deletion of *TREM2* in human microglia led to a significant reduction of the HLA transcriptional response (unpaired t-test, *p<0.05, **p<0.01). **f**, Phenotypic trajectory followed by the human microglia obtained by an unbiased pseudotime ordering with Monocle 3. Proportion of cells (y-axis) present at different stages of the pseudotime trajectory (x-axis), colored by clusters shown in Fig. 1a. Note the shift in transcriptional states in H9-*WT* (n=2) and H9-*TREM2*^*-/-*^ cells consistent with a reduction in the HLA transcriptional state. **g**, Density plots displaying the average distribution of human microglia transplanted in *App*^*hu/hu*^ mice and challenged with scrambled peptide (Scr, n=6) or soluble amyloid-β aggregates (oAβ, n=13, density is normalized by sample size). **h**, Distribution and proportion of cells across all identified clusters. Acute injection of soluble amyloid-β aggregates induced a dramatic increase in both CRM-1 and CRM-2 transcriptional profiles (each dot represents a single mouse, unpaired t-test, ***p<0.001, ****p<0.0001). **i**, Phenotypic trajectory followed by the human microglia obtained by an unbiased pseudotime ordering with *Monocle 3*. Proportion of cells from different injections (y-axis) present at different stages of the pseudotime trajectory (x-axis), colored by clusters shown in Fig. 1a. Note the shift in transcriptional states of the oAβ injected cells consistent with an increase in CRM-1 and -2 transcriptional states. **j**, Ablation of the response to amyloid-β plaques but not oligomeric Aβ in *App*^*Nl-G-F*^ mice. Correlation analysis of the logFC in microglia transplanted in *App*^*NL-G-F*^ *ApoE*^*-/-*^ vs. *App*^*NL-G-F*^ mice (y-axis) and H9-*TREM2*^*-/-*^ vs. H9-*WT* from *App*^*NL-G-F*^ mice (x-axis) (Pearson’s correlation, R= 0.17, differentially expressed genes were adjusted using Bonferroni correction and colored according to clusters in Fig. 1a). Note the common downregulation of DAM-HLA profiles, upregulation of CRMs, while IRM genes are moving mainly along the y axis. **k**, Full spectrum of amyloid-β induced responses. Correlation analysis of the logFC in microglia either challenged with oligomeric Aβ (oAβ) vs. scrambled peptide (Src, y-axis) or transplanted in *App*^*NL-G-F*^ mice (x-axis) (Pearson’s correlation, R= 0.06, differentially expressed genes adjusted using Bonferroni correction). Note how the two axes of activation are uniquely enriched in specific microglial cell states CRM versus HLA). **l**, Microglial response to oligomeric amyloid-β. Correlation analysis of the logFC in microglia either challenged with oAβ vs. Src (y-axis) or transplanted in *App*^*NL-G-F*^ *ApoE*^*-/-*^ vs. *App*^*NL-G-F*^ mice (x-axis) (Pearson’s correlation, R= 0.05, differentially expressed genes were adjusted using Bonferroni correction and colored according to clusters in Fig. 1a). Note in comparison with (k) how the CRM y axis is maintained while the DAM-HLA enrichment is reversed in absence of *ApoE*.

We compared the transcriptomic profiles of human microglia transplanted in *App*^*NL-G-F*^(n=23) and *App*^*hu/hu*^ (n=10, 3 independent cell lines per group) mice to evaluate to what extent different microglia phenotypes were induced by amyloid-β pathology. DAM, HLA, and CRM-1 clusters were significantly enriched in *App*^*NL-G-F*^ compared to *App*^*hu/hu*^ mice (Fig. 2a, b and Extended Data Fig. 5a, b). Trajectory analysis confirmed that microglia follow three main activation routes from homeostatic towards four distinct transcriptional cell states: DAM, HLA, CRM, and IRM (Fig. 2c, d and Extended Data Fig. 5c, d). Whereas the DAM to HLA and IRM trajectories partially overlap, CRM deflects early during the phenotypic transition resulting in an independent response program (Fig. 2 and Extended Data Fig. 5c, d). HLA appears as a continuation of the DAM response, resulting in a DAM to HLA trajectory. Both the presence and magnitude of these phenotypic programs are human specific features, as only small traces of HLA and CRM clusters have been observed previously in mouse models even in advanced stages of disease^3–5,10^. Expression of various genes of the HLA cluster has been associated with dense core amyloid-β plaques in human AD brains, as well as in demyelinating regions in multiple sclerosis cases^29^. We explored whether the relationship between DAM and HLA could be the equivalent to DAM1 vs. DAM2 observed in mice^4^. Direct comparison between these responses confirmed that the HLA response is unique to human microglia, and different from the murine DAM2 (Extended Data Fig. 5e). Both HLA and DAM2 showed increased expression of DAM genes (such as *CD9, CSTD* and *GPNMB*), however, HLA is characterized by an induction of antigen presenting molecules (including *HLA-DQB1, HLA-DRB5* and *CD74*), whereas DAM2 shows enrichment in ribosomal genes (like *RPS3, RPS9* and *RPS19*). This indicates that human microglia display an elaborate response to amyloid-β pathology, and actively engage in antigen presentation of phagocyted material (e.g. amyloid-β) and recruitment of the adaptive immune system. Previous findings of clonal expansion of CD8 T-cells in the brain and cerebrospinal fluid of AD patients^30^ support this hypothesis.

**Figure 5.**
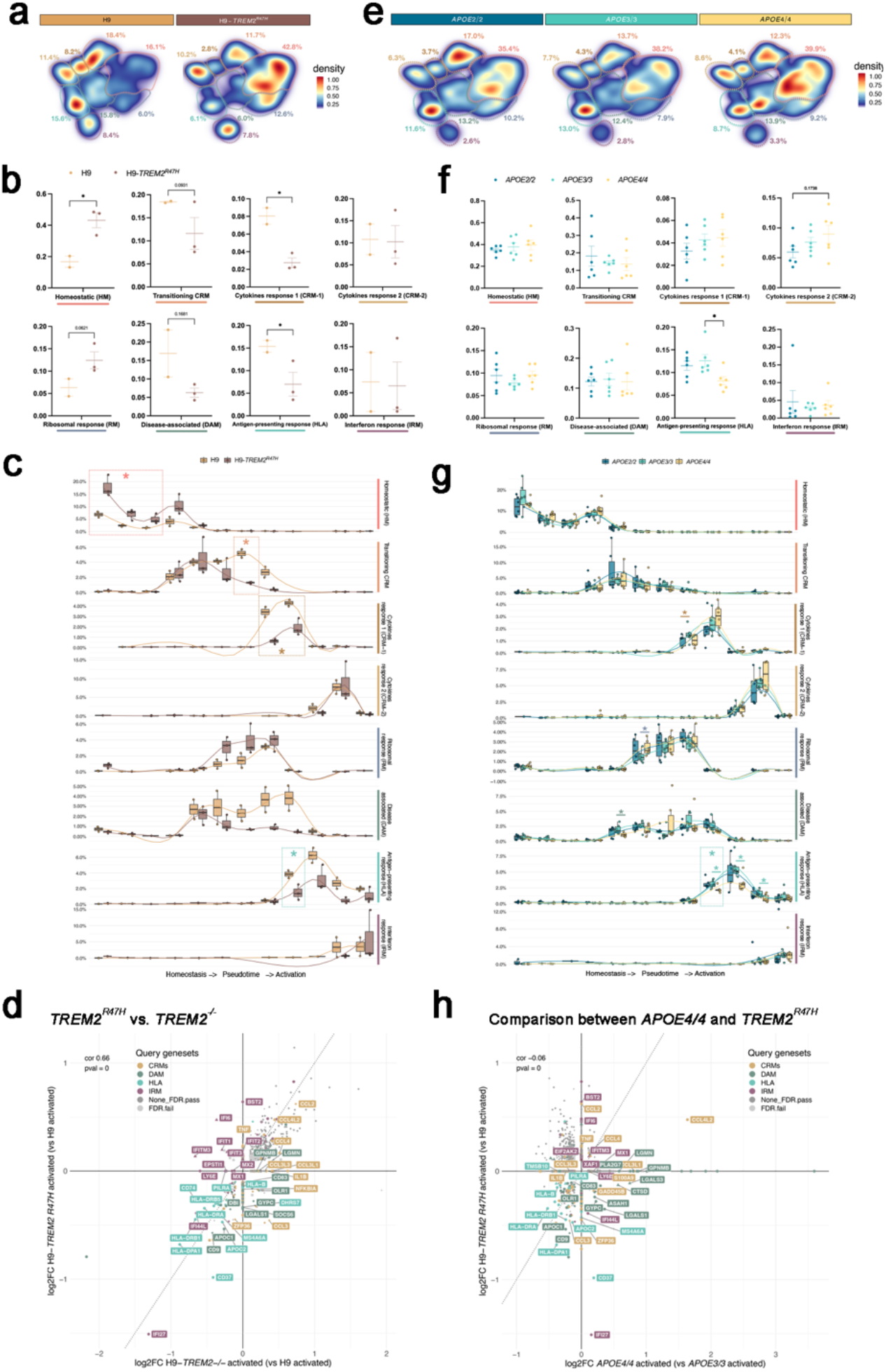
Alzheimer’s disease genetic risk modifies the response of human microglia to amyloid-β pathology. a-d Effect of the *TREM2*^*R47H*^ mutation o response to amyloid-β pathology. **a**, Density plots displaying the average distribution of human H9-*WT* (n=2) and H9-*TREM2*^*R47H*^ (n=3) microglia transplanted in *App*^*NL-G-F*^ mice. **b**, Distribution and percentage of cells across all identified clusters. *TREM2*^*R47H*^ in human microglia led to a significant increase in HM and reduction of HLA and CRM-1 transcriptional responses (unpaired t-test, *p<0.05). **c**, Phenotypic trajectory followed by the human microglia obtained by an unbiased pseudotime ordering with Monocle 3. Proportion of cells (y-axis) present at different stages of the pseudotime trajectory (x-axis), colored by genotypes shown in **a**. The data is displayed as box plots, showing all individual replicates for each pseudotime bin. Note the strong shift in transcriptional states in H9-*TREM2*^*R47H*^ cells consistent with a reduction in HLA and CRM transcriptional states. **d**, *TREM2*^*-/-*^ vs. *TREM2*^*R47H*^. Correlation analysis of the logFC in *TREM2*^*R47H*^ vs. H9-*WT* (y-axis) and H9-*TREM2*^*-/-*^ vs. H9-*WT* (x-axis) microglia transplanted in *App*^*NL-G-F*^ mice (Pearson’s correlation, R= 0.17, differentially expressed genes were adjusted using Bonferroni correction and colored according to clusters in Fig. 1a. **e-h** Effect of the different *APOE* allelic variants in the response of microglia to amyloid-β pathology. **e**, Density plots displaying the average distribution of human microglia harboring the *APOE2/2* (n=6), *APOE3/3* (n=6) and *APO4/4* allelic variants and transplanted in *App*^*NL-G-F*^ mice. **f**, Distribution and percentage of cells across all identified clusters. *APOE4/4* in human microglia led to a significant reduction of the HLA transcriptional response compared to *APOE3/3* cells (unpaired t-test, *p<0.05). **e**, Phenotypic trajectory followed by the human microglia obtained by an unbiased pseudotime ordering with Monocle 3. Proportion of cells (y-axis) present at different stages of the pseudotime trajectory (x-axis), colored by genotypes shown in **b**. The data is displayed as box plots, showing all individual replicates for each pseudotime bin. Note the shift in transcriptional states in *APOE4/4* cells consistent with a reduction in the HLA transcriptional state. **h**, Comparison of *TREM2*^*R47H*^ and *APOE4* effects. Correlation analysis of the logFC in *TREM2*^*R47H*^ vs. H9-*WT* (y-axis) and *APOE4/4* vs. *APOE3/3* (x-axis) microglia transplanted in *App*^*NL-G-F*^ mice (Pearson’s correlation, R= 0.17, differentially expressed genes were adjusted using Bonferroni correction and colored according to clusters in Fig. 1a). Note the downregulation of HLA genes in both clinical mutations increasing AD risk.

Overall human microglia display a complex, multi-pronged response to amyloid-β pathology, including some programs that share features of the previously described DAM, IRM and TRM responses^3,4^, but also two human specific CRM and HLA transcriptional states. This defines a heterogeneous transcriptional landscape in response to amyloid-β pathology that we propose to group under the term human Amyloid-β Response Microglia or (human) ARM.

### Human microglial responses in post-mortem tissue

We extracted microglial single nuclei RNA-sequencing data from three independent studies investigating the transcriptome of human microglia in the AD brain: 176,136 nuclei from Gerrits et al. ^21^; 28,767 from Sayed et al^31^; and 3,978 from Zhou et al.^19^. We reproduced their original clustering analysis, and generally found that there is no congruency between these post-mortem datasets (Extended Data Fig. 6). We re-analyzed these data using our transcriptomic profiles from transplanted microglia (Fig. 3a-c). Whereas the clusters reported across these studies showed little overlap, we were able to identify all our microglial cell states in the three data sets, especially HM, DAM, HLA and CRM (Fig. 3 and Extended Data Fig. 7). The IRM signature was more elusive, but still overlapping with clusters in two of the three studies, and confirmed in another single cell study using freshly isolated microglia from surgical resections^32^. Importantly, none of the three post-mortem datasets alone covered the full phenotypic diversity in response to amyloid-β as observed in the xenotransplanted microglia (Fig. 3d). For example, in Gerrits *et al*.^21^ DAM corresponds to *AD1*, and CRM to *Inflammation*, but the HLA signature is absent (Fig. 3a,d). In Zhou et al. ^19^, all signatures are present, but they converge in a limited number of clusters (*Micro1, Micro4* and *Micro3*). In Sayed *et al*. ^31^ although the main amyloid-β induced signatures are present, DAM and HLA, and IRM and CRM show partial overlap (Fig. 3b,d). Certain transcriptomic states from primary samples which were correlated with tau pathology (e.g. *AD2*) overlap with our HM signature. Additional cell states described in these post-mortem studies but not captured in our xenotransplantation model might reflect the response to additional pathologies in the late phase of AD (Fig. 3d and Extended Data Fig. 7b). Overall, we not only provide solid evidence that the cell states in transplanted microglia are present in the human AD brain, but also demonstrate that re-analysis and re-interpretation of available human post-mortem datasets using single cell profiles from xenotransplanted human microglia as a reference increases the consistency across studies and allows to discriminate microglia responses to amyloid-β pathology from responses to Tau- and other pathologies.

**Figure 6.**
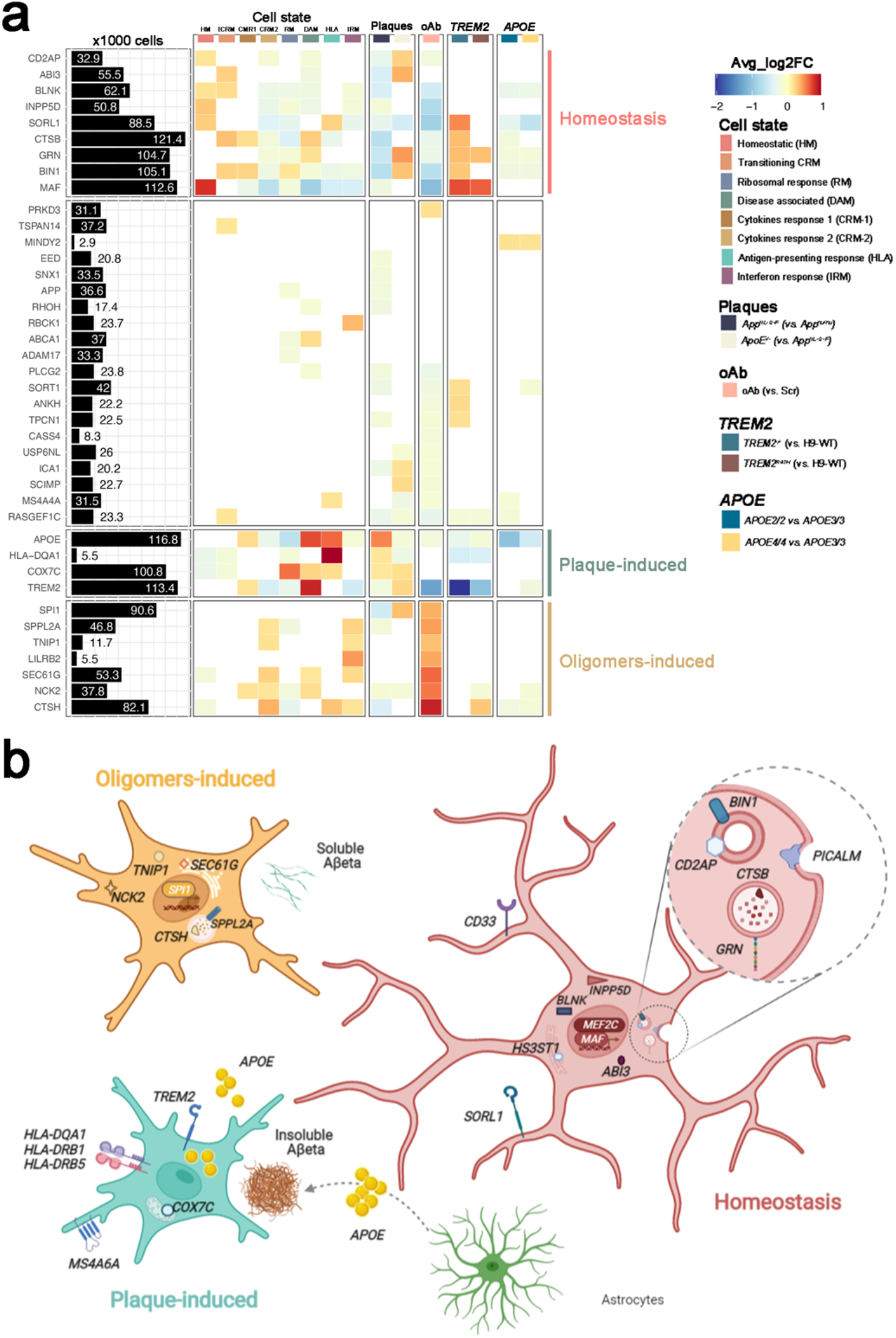
Alzheimer’s disease genetic risk genes are differentially expressed in human microglial cell states and modulated by amyloid-β pathology. **a**, Analysis of GWAS genes enrichment in xenotransplanted microglia. The black bars represent the number of cells (in thousands) with detectable expression (>=1 read per cell) for each candidate gene. The heatmap summarizes the deregulated expression (LogFC, colour scale) of these genes across cell states (each cluster compared to all others), as well as after exposure to amyloid-β plaque pathology, upon injection of soluble oligomeric Aβ, or altering the genetic background of the mice or the transplanted cells. The genes are ranked in rows based on hierarchical clustering. We identify 3 sets of genes that display a common profile across cell states (based on their enrichment in the specific microglial phenotypic transcriptional states HM, DAM and HLA, and CRM), amyloid-β pathology and genetic risk, and we group these profiles as: microglia Homeostasis, plaque-induced genes, and oligomers-induced genes. The remaining genes did not show a clear enrichment in cell states or other conditions. All differential expressions were significant after adjusting P-values using Bonferroni correction (FDR < 0.05). Only genes that are significantly changing in at least one of the tested conditions are reported, see Supplementary Table 7 and Extended Data Fig. 12 for further details. **b**, Illustration of the complex microglial AD genetics by cell-states profiles and driven by different amyloid-β pathologies as found in (a) and Extended Data Fig. 12.

### Human microglia engage in different activation programs in response to distinct forms of amyloid-β pathology

We wondered whether the distinct transcriptional trajectories adopted by human microglia might be caused by different forms of amyloid-β pathology. We used a genetic approach to reduce the amyloid-β plaques in the mouse brain by knocking out *ApoE* (*App*^*NL-G-F*^ *ApoE*^*-/-*^). As previously reported, this leads to a significant reduction of amyloid-β plaques^33^ (Extended Data Fig. 8). We analyzed 22,387 microglia isolated from 6 months old *App*^*NL-G-F*^ (n=11) and *App*^*NL-G-F*^ *ApoE*^*-/-*^ (n=6, 2 independent iPSC lines per group) (Extended Data Fig. 2 and 3). Clustering analysis revealed a significant reduction in the recruitment of microglia into DAM and HLA phenotypes, with no alterations in the CRM response (Fig. 4a, b). Trajectory analysis confirmed a strong reduction in the proportion of cells transitioning into the DAM-HLA transcriptional axis (Fig. 4c and Extended data Fig. 9). This also indicates that the main impact of *ApoE* on microglia is indirect by the modulation of amyloid-β deposition, rather than a direct, cell-autonomous effect on their transcriptomic response. To confirm this, we investigated whether *TREM2*, the main cell-autonomous mediator of the microglia response to amyloid-β plaques in mice^4,5,34^, was able to modify human microglial cell states in a similar way. Analysis of 3,282 H9 (n=2) and 1301 H9-*TREM2*^*-/-*^ (n=3) microglia from 6 months old *App*^*NL-G-F*^ mice confirmed the suppression of DAM and HLA responses, but again without altering the CRM profile (Fig. 4d-f, l and Extended Data Fig. 9e-h). Thus, DAM and HLA responses are dependent on amyloid-β plaques and mediated by TREM2. Contrary to mouse systems, human cells do not display the DAM1/DAM2 duality^4^. Our data shows that TREM2 mediates the phenotypic transition to both DAM and HLA highlighting species-specific differences in how microglia react against the fibrillar amyloid-β present in plaques.

To further confirm that the remaining CRM response is mediated by soluble amyloid-β, we injected synthetic amyloid-β oligomers (5μl at 10μM, n=13) or scrambled peptide (Scr, n=7, 2 independent cell lines per group) in the ventricle of xenotransplanted 3 months old *App*^*hu/hu*^ mice^15^. We isolated and sequenced 29,658 microglia 6h after injection (Extended Data Fig.2). Whereas microglia exposed to Scr peptides remain largely homeostatic, almost all microglia challenged with soluble amyloid-β oligomers adopt a CRM transcriptional state (Fig. 4g, h and Extended Data Fig. 9i-l). Trajectory analysis confirmed that soluble amyloid-β treated cells undergo an almost complete phenotypic transition across CRM-1 and CRM-2 clusters (Fig. 4i). CRM-2 is at the extreme of this trajectory and remains unchanged in *App*^*NL-G-F*^ mice but becomes the dominant cell state after acute injection of high levels of amyloid-β oligomers (Fig. 4g-I, j, k), consistent with our previous observations^22^. Thus, overall, the ARM response consists of independent cell states that co-exist in the *App*^*N-L-GF*^ model for AD and are elicited by amyloid-β plaques (DAM and HLA) and soluble amyloid-β species (CRM), respectively.

### Alzheimer’s disease risk genes modulate human microglial cell states in vivo

Our model is very well suited to investigate how AD genetic risk modulates human microglia responses. As proof-of-concept, we investigated some of the major genetic risk factors for AD, hypothesizing that they would modulate microglia according to their role in AD risk, i.e. *APOE2/2* will increase protective and/or decrease damaging responses, whereas *APOE4/4* and *TREM2* mutations will increase damaging or reduce protective responses^1^. We first investigated the effect of the clinical mutation *TREM2*^*R47H*^ generated on the H9 background (Table 1). We profiled 3,282 H9 (n=2) and 5,845 H9-*TREM2*^*R47H*^ (n=3) microglia transplanted in *App*^*NL-G-F*^ mice (Extended Data Fig. 2 and 3). The *TREM2*^*R47H*^ mutation induced a significant reduction in the fraction of cells recruited into HLA, but also an unexpected reduction of the CRM-1 cluster (Fig. 5a, b and Extended Data Fig. 10a-d). Trajectory analysis confirmed that this clinical mutant remains largely locked in a homeostatic state (Fig. 5c and Extended Data Fig.10c, d). We compared the gene expression alterations induced by complete loss-of-function of *TREM2* and the clinical *TREM2*^*R47H*^ mutant (Fig. 5d). Both genotypes show a high correlation and a common downregulation of DAM and HLA genes. However, whereas TREM2^−/-^ retains expression of CRM genes (including *IL1B, CCL3L3 or CCL3L1*), *TREM2*^*R47H*^ shows spared expression of IRM genes (such as *IFIT1, IFIT3, IFI6* or *IFITM3*). Our data indicate that the *TREM2*^*R47H*^ results in an inability of microglia to engage into both HLA and CRM transcriptional programs, and therefore do not react against amyloid-β plaques and oligomers, respectively.

Upregulation of *ApoE* in microglia is one of the main transcriptional changes induced by amyloid-β plaques in mouse systems^4,5,7^. This suggests that the *APOE* allelic variants could be important regulators of human microglia. We used a series of isogenic *APOE2/2, 3/3* and *4/4* iPSC lines (UKBIO11-A, EBISC). We also obtained a second series of the *APOE* alleles in an *APOE* knock out iPSC line (BIONi010-C, EBISC) with *APOE2/0, 3/0* and *4/0* genotypes (Table 1, Extended Data Fig. 2 and 3). Both series have very similar *APOE* expression levels (Extended Data Fig. 11) and we therefore grouped them together for our analyses. We transplanted microglia precursors from all these lines into *App*^*NL-G-F*^ mice (n=3 per cell line). Unlike *TREM2*^*R47H*^ cells, clustering analysis did not reveal major differences between cells harboring the different *APOE* allelic variants. *APOE4/4* expressing microglia show, however, a significant reduction in the proportion of cells acquiring the HLA phenotype and a tendency towards an increase in CRM response (p=0.17, Fig. 4e,f and Extended Data Fig. 10e, f). Trajectory analysis confirmed this and revealed that *APOE4* cells accumulate in the last stages of DAM (Fig. 5g and Extended Data Fig. 10g, h), suggesting an inability to transition from DAM towards HLA, and therefore to a full response to Aβ plaques. Despite *APOE4* being the major genetic risk factor for AD, our data suggest that its cell-autonomous role in microglia is rather limited, probably reflecting the fact that it operates across multiple cell types as recently shown for example for astrocytes and microglia^35,36^. As shown above, the main effect of *ApoE* knock out models on microglia responses seems non-cell autonomous via the decreased accumulation of amyloid-β plaques. Nevertheless, the changes induced by *TREM2*^*R47H*^ and *APOE4* show a common impairment in activating the human specific HLA (rather than DAM) transcriptional program and thus suggest that the HLA response to amyloid-β plaques might be a beneficial aspect of the response of human microglia to AD pathology (Fig. 5h).

Finally, we extracted a list of 85 GWAS significant gene candidates (nearest gene to significant genetic loci, P-value < 5×10^−8^) from the most recent meta-analysis of AD complex genetics^37^. We found that almost 90% of these genes were detected in our dataset (Supplementary Table 7 and Extended Data Fig. 12a) while more than half were differentially expressed in the conditions we tested, confirming that a large part of the genetic risk for AD is harbored by microglia and that their expression is modified in the context of amyloid-β pathology^6^ (Fig. 6a and Extended Data Fig. 12). We observed that different subsets of AD risk genes were enriched in all cell states in response to amyloid-β pathology. Whereas genes like *ABI3, INPP5D, SORL1, GRN, BIN1* and *MAF* were enriched in homeostatic microglia, both the response to amyloid-β plaques and oligomers were associated with distinctive groups of genes including *TREM2, APOE, COX7C* and *HLA-DQA1* in DAM/HLA; and *SPl1, SPPL2A, TNIP1, LILRB2, SEC61G, NCK2* and *CTSH* in the response to amyloid-β oligomers (Fig. 6a and Extended Data Fig. 12). We also observed altered expression of subsets of these AD-risk genes after introducing clinical mutations in both *APOE* and TREM2. Consistent with their inability to transition towards responsive cell states, both *TREM2*^*-/-*^ and *TREM2*^*R47H*^ microglia were enriched compared to H9-WT in those risk genes present in homeostatic cells, such as *SORL1, GRN, CSTB, BIN1* and *MAF*; and reduced expression of DAM and HLA genes like *APOE* or *HLA-DQA1* (Fig. 6a). *APOE2/2* and *APOE4/4* microglia showed milder risk genes enrichment when compared to *APOE3/3*, but in a similar direction with decreased association to DAM and HLA genes (Fig. 6a and Extended Data Fig. 12). Our data show that the contribution of AD genetics to the biology of microglia is complex, likely having an impact on multiple cell states. It is interesting to see how known genetic risk factors affect the expression of other genetic risk factors in microglia, providing a strong basis for the concept of polygenic risk and providing a frame to understand the functional implications of polygenic risk (Fig. 6b).

Overall, microglia respond to AD’s multi-faceted Aβ pathology, with a complex, multipronged human Amyloid-β Response Microglia or (human) ARM. Unique features appear the HLA response and the strong CRM response to oligomeric amyloid-β which are not recapitulated in mouse systems. We also demonstrate that genetic risk of AD significantly alters the response of human microglia to amyloid-β pathology. The blocking effects of *APOE4* polymorphism or the *TREM2*^*R47H*^ mutation on the HLA response to amyloid-β plaques, which has no clear mouse counterpart, suggest that this aspect of the microglial response is protective against developing AD. On the other hand, it is unclear how the CRM response against soluble amyloid-β species contributes to AD pathology. CRM is characterized by the upregulation of classical inflammatory cytokines and chemokines such as *IL1B, CCL2, CCL3, CCL4* and components of the *NFKB* pathway. It is also enriched in several AD risk genes. Soluble amyloid-β species appear early during the disease process long before amyloid-β plaques and have been linked to early neuronal dysfunction^38^. Our data suggests that microglia might also engage in AD at very early disease stages. The question remains as to whether the upregulated cytokines and chemokines characteristic for this response affect neurons or other brain cells, inducing the cellular responses in AD that ultimately result in neurodegeneration^39^.

Our findings unravel the complex roles of human microglia in early and late amyloid-β pathology of AD. These results indicate that therapeutic targeting of microglia needs to be implemented with care as it might differentially affect their cell states and modify the disease course in unpredictable ways. Our data provide an invaluable foundation to further explore the multi-pronged contribution of human microglia to AD.

## Authors contribution

R.M., N.F, A.M.M. and B.D.S. conceived and designed the study and wrote the manuscript. R.M, N.F and A.M.M. performed experiments and analyzed data together with B.D.S. E.D. provided bioinformatic support and performed the analyses on publicly available datasets. L.W. performed experiments. J.V.D.D. performed the experiments involving *TREM2* mutant microglia. I.G. performed mouse brain surgery and injections of amyloid-beta oligomers in the brain. P.P. assisted on the experiments involving *ApoE*^*-/-*^ mice. L.S. generated the *App*^*hu/hu*^ and *ApoE*^*-/-*^ mouse lines. S.P. assisted in the generation of cDNA libraries for sequencing. S.B. generated and provided the iCas9-H9 stem cell line. C.V. provided supervision to J.V.D.D. M.F. provided supervision to E.D. All authors discussed the results and commented on the manuscript

## Acknowledgments

This project received funding from the European Research Council (ERC) under the European Union’s Horizon 2020 Research and Innovation Programme (grant agreement no. ERC-834682 CELLPHASE_AD). This work was also supported by UK Dementia Research Institute, the Flanders Institute for Biotechnology (VIB vzw), Vlaams Initiatief voor Netwerken voor Dementie Onderzoek (Strategic Basic Research Grant no. 135043), a Methusalem grant from KU Leuven and the Flemish Government, the Fonds voor Wetenschappelijk Onderzoek, KU Leuven, The Queen Elisabeth Medical Foundation for Neurosciences, the Opening the Future campaign of the Leuven Universitair Fonds, the Belgian Alzheimer Research Foundation (SAO-FRA) and the Alzheimer’s Association USA. R.M. has funding from Fonds voor Wetenschappelijk Onderzoek (grants no. G0C9219N, G056022N and G0K9422N) and is a recipient of a postdoctoral fellowship from the Alzheimer’s Association USA (fellowship no. 2018-AARF-591110 and 2018-AARF-591110-RAPID). He also receives funding from BrightFocus Foundation (A2021034S), SAO-FRA (grant no. 2021/0021) and the University of Antwerp (BOF-TOP 2022-2025). B.D.S. holds the Bax-Vanluffelen Chair for Alzheimer’s Disease. B.D.S. receives funding from the Medical Research Council, the Alzheimer’s Society and Alzheimer’s Research UK via the Dementia Research Institute. N.F. is recipient of a PhD fellowship from Fonds voor Wetenschappelijk Onderzoek (fellowship no. 1139520N). A.M.-M. is supported by a fellowship from the Alzheimer’s Association USA (fellowship no. AARF-20-684397) and a Marie Skłodowska-Curie Actions - Seal of Excellence Postdoctoral Fellowship (12ZX621N). We thank V. Hendrickx and J. Verwaeren for animal husbandry. Imaging was acquired through Nikon A1R Eclipse Ti confocal from LiMoNe facility at CBD.

## Methods

### Mice

This *App* single knock-in mouse model (*App*^*NL-G-F*^; Takaomi Saido)^40^ does not overexpress APP like classical APP mouse models, but contains the humanized Aβ sequence, as well as Swedish (NL), Arctic (G), and Iberian (F) mutations. *App*^*NL-G-F*^ mice accumulate amyloid-β plaques and suffer from learning, memory, and attention impairments from 6 months onwards^40,41^. The humanized *App*^*hu/hu*^ mice have been recently generated in our lab to serve as controls^28^. Both strains were crossed with homozygous *Rag2*^*tm1*.*1Flv*^ *Csf1*^*tm1(CSF1)Flv*^ *Il2rg*^*tm1*.*1Flv*^ *App*^*tm3*.*1Tcs*^ mice (Jacksons Lab, strain 017708) to generate the *Rag2*^*-/-*^ *Il2rγ*^*-/-*^ *hCSF1*^*KI*^ *App*^*NL-G-F*^ and the *Rag2*^*-/-*^ *Il2rγ*^*-/-*^ *hCSF1*^*KI*^ *App*^*hu/hu*^ used in this study. In total, we transplanted >125,000 cells across 11 different cell lines, and three mouse background genotypes. For all experiments, we used Total-Seq A hashing antibodies (Biolegend) to be able to demultiplex individual mouse replicates (Tables S1, S3 and Extended Data Fig. 2). Mice had access to food and water *ad libitum* and were housed with a 14/10 h light-dark cycle at 21°C in groups of two to five animals. All experiments were conducted according to protocols approved by the local Ethical Committee of Laboratory Animals of the KU Leuven (ECD project number P177/2017) following country and European Union guidelines.

### Generation of Rag2^-/-^ Il2rγ^-/-^ hCSF1^KI^ App^NL-G-F^ ApoE^-/-^ mice

*ApoE* knock out mice were generated in zygotes from homozygous *Rag2tm1*.*1Flv Csf1tm1(CSF1)Flv Il2rgtm1*.*1Flv Apptm3*.*1Tcs* mice using CRISPR-Cas9 technology by targeting exon4 of the mouse *ApoE* gene. The RNA guide 5′-CCTCGTTGCGGTACTGCCCGAGT-3 was selected using the CRISPOR web tool. Ribonucleoproteins (RNPs) containing 0.3 μM purified Cas9HiFi protein (Integrated DNA Technologies, IDT), 0.3 μM CRISPR RNAcrRNA and 0.3 μM trans activating crRNA (IDT) were injected into the pronucleus of 72 embryos by microinjection in the Mouse Expertise Unit of KU Leuven. Two candidate pups were identified by PCR analysis with several primer combinations. One founder was selected for breeding and an allele with a chromosomal deletion of 335 bp (corresponding to 148 bp of intronic sequence and the first 187 bp of exon 4 sequence (Extended Data Fig. 8) was selected to establish the colony. The founder mouse was backcrossed over two generations before a homozygous colony was established, which was designated *App*^*NL-G-F*^ *ApoE*^*-/-*^. The strain is maintained on the original C57Bl6J:BalbC background. Standard genotyping for the *ApoE* allele is performed by PCR with primers 5′-GCTCCCAAGTCACACAAGAA-3’ and 5’-CTCACGGATGGCACTCAC-3’ resulting in a 755 bp amplicon for the wildtype allele and a 420 bp amplicon for the *ApoE* knock-out allele.

### Microglial progenitors differentiation

UKBIO11-A, BIONi010-C, H9-WA09 and their isogenic modifications (Table 1) were differentiated into microglial precursors and transplanted following our recent published protocol, MIGRATE^23^. In brief, stem cells were plated and maintained in human matrigel coated 6-well plates and in E8 flex media until reaching ∼70-80% confluence. Once confluent, stem cell colonies were dissociated into single cells and plated into U-bottom 96 well plates at a density of ∼10k/well in mTeSR1 media with BMP4 (50 ng/ml), VEGF (50 ng/ml) and SCF (20 ng/ml) for 4 days. On day 4, embryoid bodies (EBs) were transferred into 6 well-plates (∼20 EBs/well) in X-VIVO (+supplements) media supplemented with SCF (50 ng/ml), M-CSF (50 ng/ml), IL-3 (50 ng/ml), FLT3 (50 ng/ml) and TPO (5 ng/ml) for 7 days with a full change of media on day 8. On day 11, differentiation media was replaced with X-VIVO (+supplements) with FLT-3 (50 ng/ml), M-CSF (50 ng/ml) and GM-CSF (25 ng/ml). On day 18, human microglial precursors were harvested and engrafted into P4 mouse brains (0.5 M cells/pup) as previously described. Prior to transplantation, mouse microglia were depleted by inhibiting CSF1 receptor (CSF1R) with BLZ945 (dose of 200 mg/kg) at P2 and P3 as previously described^15,23^.

### Genetic modification of stem cell lines

Generation of *TREM2*^*-/*−^ and *TREM2*^*+/R47H*^ from H9 wild-type (WT, WA09) human embryonic stem cells was done as described in Claes et al., (2019). Briefly, for the *TREM2*^*+/R47H*^ CRISPR/Cas9 nickases and two guide RNAs (gRNA A and B) that target exon 2 of *TREM2* nearby the location of *R47H* (*G>A*) and a genomic *TTAA* were purchased from Addgene. A donor plasmid was made comprising homology arm 1 (HA1) of *TREM2* (with the *R47H* mutation), a selection cassette (CAGG promoter, HYG/TK, green fluorescent protein) and HA2 of *TREM2* exon 2. To create *TREM2*^−/−^ hPSCs, a CRISPR/Cas9, gRNA B, and the same donor plasmid were used. To create *TREM2*^*+/R47H*^ hPSCs, 2 × 10^6^ single cells of the heterozygously targeted clone were nucleofected with 4 μg of piggyBac (PB) transposase plasmid and negative selection with fialuridine, also known as 1-(2-deoxy-2-fluoro-1-D-arabinofuranosyl)-5-iodouracil (FIAU) (1:8000– 1:2500; 0.5 mM in water), was applied to select for cells wherein the selection cassette was removed. Of note, the H9 WT line from which the *TREM2*^*+/R47H*^ and *TREM2*^−*/*−^ lines were created carries an APOE ε3/ε4 genotype.

To get the gRNA into the cells, nucleofection was performed. Briefly, 2×10^6 single cell H9s were pre-incubated with Revitacell® (Life Technologies) and nucleofected using the Amaxa nucleofector II on setting F16 with 2.5μg CRISPR/Cas9, 2.5 μg gRNA (A and) B and 5μg donor template to create TREM2-/-hPSCs. Selection was initiated after 2-3 days with 25-150μg/ml of Hygromycin B (Sigma-Aldrich) and maintained for 10-15 days. Recombinant colonies were manually picked and expanded for further characterization.

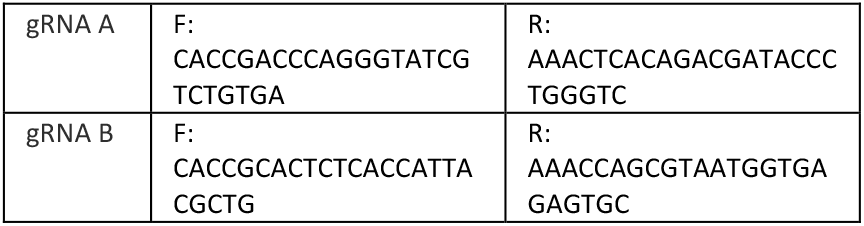

H9 (WA-09) embryonic stem cell line were also modified to express the CAS9 in doxycycline dependent manner (iCas9), using TALENS at the AAVS1 locus to facilitate the generation of targeted gene deletion. H9-iCas9 were only used in libraries 14 and 16 and doxycycline was never administered.

### Apoe expression using qPCR

The levels of *Apoe* expression in the grafted cell lines were checked by collecting cell pellets. Using the RNeasy Micro Kit, RNA was extracted according to manufacturer’s instructions and the RNA was reverse transcribed with the High-Capacity cDNA Reverse Transcription Kit. A qPCR was performed with SensiFast SYBR reagent and custom-made primers for GAPDH (FW: tcaagaaggtggtgaagcagg; RV: accaggaaatgagcttgacaaa) and Apoe (we average the level of expression of multiple primers spanning the whole gene, see table below).

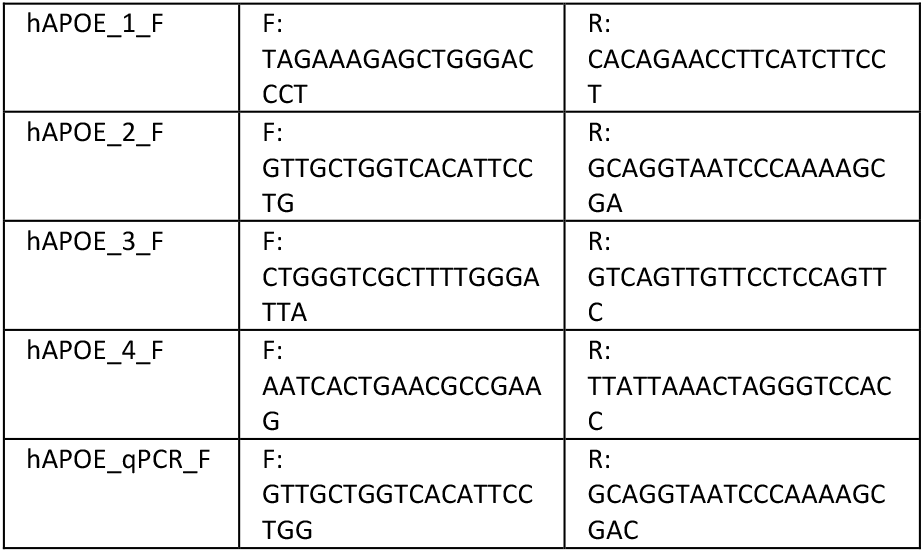

### Soluble amyloid-β preparation and ICV injections

Soluble amyloid-β aggregates (10 μM) or scrambled (10 μM) were prepared as previously ^15,42^. Briefly, recombinant amyloid-β (1-42) or scrambled were thawed during 30 minutes at room temperature and dissolved in hexafluoroisopropanol (HFIP) at 2mg/ml. HFIP was fully evaporated with a gentle stream of N2 gas and resulting peptides were dissolved in dimethyl sulfoxide (DMSO) at 2mg/ml. DMSO media was removed using HiTrap® Desalting column 5kD and peptides were eluted in Tris-EDTA buffer. Of note, Tris-EDTA buffer was composed of 50 mM Tris-buffer, 1 mM EDTA at pH= 7.5. Tris-EDTA-eluted amyloid-β or scrambled peptides were quantified using Bradford assay prior to oligomerization. Peptides were let oligomerize for 2h at room temperature. After 2h, Amyloid-β (1–42) or scrambled oligomers were diluted to a final concentration of 10 μM in Tris-EDTA buffer, snap frozen and stored at -80°C. Following a similar approach as previously described^15^, at 12 weeks of age, *App*^*hu/hu*^ mice engrafted with the full isogenic series of UKBIO11-A or BIONi010-C were anaesthetised with isoflurane and injected intracerebroventricularly (ICV) with 5 μl of oligomeric Aβ (10 μM) or scrambled peptide (10 μM). Stereotactic coordinates from Bregma: anteroposterior: - 0.22 mm; mediolateral: -1 mm; dorsoventral: -2.74 mm. After surgery, mice were placed on a thermal pad until recovery. 6 hours after injection, *App*^*hu/hu*^ mice were euthanized, and human microglia were isolated using FACS sorting for transcriptomics analysis.

### Human microglia isolation from mouse brain for single-cell transcriptomics

At 6-7 months of age *App*^*NL-G-F*^, *App*^*NL-G-F*^ *ApoE*^*-/-*^ and *App*^*hu/hu*^ xenotransplanted with H9, UKBIO11-A, BIONi010-C (Table 1) and their isogenic modifications were sacrificed with an overdose of pentobarbital and immediately perfused with ice-cold 1x DPBS (Gibco, Cat#14190-144) supplemented with 5U of heparin (LEO). After perfusion, 1 hemisphere of each mouse brain without cerebellum and olfactory bulbs was placed in FACS buffer (1x DPBS, 2% FCS and 2 mM EDTA) + 5 μM Actinomycin D (ActD, Sigma, Cat#A1410-5MG) for transcriptomics. Brains were mechanically and enzymatically dissociated using Miltenyi neural tissue dissociation kit P (Miltenyi, Cat#130-092-628) supplemented with 5 μM ActD. Next, samples were passed through a 70 μm strainer (BD2 Falcon), washed in 10 ml of ice-cold FACS buffer + 5 μM ActD and spun at 300g for 15 minutes at 4°C. Note that 5 μM ActD was kept during collection and enzymatic dissociation of the tissue to prevent artificial activation of human microglia during the procedure as previously reported^13^. ActD was removed from the myelin removal step to prevent toxicity derived from long-term exposure. Following dissociation, myelin was removed by resuspending pelleted cells into 30% isotonic Percoll (GE Healthcare, Cat#17-5445-02) and centrifuging at 300g for 15 minutes at 4°C. Accumulating layers of myelin and cellular debris were discarded and Fc receptors were blocked in FcR blocking solution (1:10, Miltenyi, Cat#130-092-575) in cold FACS buffer for 10 minutes at 4°C. Next, cells were washed in 5 ml of FACS buffer and pelleted cells were incubated with the following antibodies: PE-Pan-CD11b (1:50, Miltenyi, Cat#130-113-806), BV421-mCD45 (1:500, BD Biosciences, Cat#563890), APC-hCD45 (1:50, BD Biosciences, Cat#555485), TotalSeq™-A cell hashing antibodies (1:500, Biolegend) and viability dye (eFluor 780, Thermo Fisher Scientific, Cat#65-0865-14) in cold FACS buffer during 30 minutes at 4°C. After incubation, cells were washed, and the pellet was resuspended in 500 μl of FACS buffer and passed through 35 μm strainer prior sorting. For sorting, cell suspension was loaded into the input chamber of MACSQuant Tyto Cartridge and human cells were sorted based on CD11b and hCD45 expression at 4 °C (Extended Data Figure 1). FACS data was analysed using FCS express software.

### Histology

When sacrificing and harvesting mouse brains for single cell sequencing, 1 hemisphere of *App*^*NL-G-F*^, *App*^*NL-G-F*^ *ApoE*^*-/-*^ *and App*^*hu/hu*^ xenotransplanted with H9, UKBIO11-A, BIONi010-C was also preserved and postfixed in 4% PFA overnight at 4°C. After 24h, PFA was removed, washed, and kept in 1x DPBS at 4°C until further processing. For sectioning, olfactory bulb and cerebellum were discarded and brains were cut coronally (40 µM thickness) with a vibrating microtome (Leica). Each sample was collected under free-floating conditions in a series of six sections and stored in cryoprotectant solution (40% PBS, 30% ethylene glycol, 30% glycerol) at -20°C. For staining, sections are washed in 1x DPBS and permeabilized for 15min at room temperature in PBS with 0.2% Triton. After permeabilization, sections were stained with X34 staining solution (10µM X34 (Sigma-Aldrich), 20mM NaOH (Sigma-Aldrich) and 40% ethanol) for 20min at room temperature. Sections were washed several times with 40% ethanol for 2min and with PBS+0.2%Triton for 5min. Finally, sections were mounted with Mowiol mounting media (Sigma-Aldrich). Pictures at 4x and 20x were taken on a Nikon A1R Eclipse confocal. Plaque analysis of the sections was quantified with ImageJ software after determining a threshold for background correction.

### Single-cell libraries preparation and sequencing

For single-cell RNA sequencing, 15,000-20,000 human microglia (CD11b+, hCD45+) from each mouse were sorted on the MACSQuant Tyto (Extended Data Fig. 1) and diluted to a final concentration of 1000 cells/µl. Since all the samples were individually hashed using TotalSeq™-A cell hashing antibodies, 2000 human microglia/animal were pooled and loaded onto the Chromium Next GEM Chip G (PN #2000177). The DNA library preparations were generated following manufacturers’ instructions (CG000204 Chromium Next GEM Single Cell 3’ Reagent Kits v3.1). In parallel the hashtag oligo libraries were prepared according to manufacturers’ instructions (BioLegend - TotalSeq™-A Antibodies and Cell Hashing with 10x Single Cell 3’ Reagent Kit v3 3.1 Protocol) using 16cycles for the index PCR. A total of 20 libraries containing 95 biological replicates were sequenced targeting 90% mRNA and 10% hashtag oligo library (50,000 reads/cell) on a HiSeq4000 or NovaSeq6000 (Illumina) platform with the recommended read lengths by 10X Genomics workflow.

### Statistics

Statistical analysis on the distribution of different experimental groups across clusters was performed using each mouse as a single replicate. We used both t-test and one-way ANOVA when comparing two or more than two groups, respectively. Statistical significance was set at a *P-value*<0.05.

### Analysis of single-cell RNA sequencing datasets

#### Alignment and Software

The raw BCL files were demultiplexed and aligned by Cellranger (v.3.1.0) against the human genome database (hg19, Ensembl 87). Raw count matrices were imported in R (v4.1.3) for data analysis. Datasets were analyzed using the *Seurat* R package pipeline (v.4.0.1) The other packages and their versions used for the analyses of this study are reported as the result of *SessionInfo()* below.

#### Attached base packages

stats4, parallel grid, stats, graphics grDevices utils, datasets methods base

#### Other attached packages

UCell_1.99.1, monocle3_0.2.3.0, SingleCellExperiment_1.12.0, SummarizedExperiment_1.20.0, GenomicRanges_1.42.0, GenomeInfoDb_1.26.4, IRanges_2.24.1, S4Vectors_0.28.1, MatrixGenerics_1.2.1, matrixStats_0.58.0, Biobase_2.50.0, BiocGenerics_0.36.0, Matrix_1.3-2, Seurat_4.0.1, wesanderson_0.3.6, VennDiagram_1.6.20, futile.logger_1.4.3, RColorBrewer_1.1-2, ggrepel_0.9.1, ggplot2_3.3.5, multcomp_1.4-18, TH.data_1.1-0, MASS_7.3-53.1, survival_3.2-13, mvtnorm_1.1-1, broom_0.7.5, magrittr_2.0.2, purrr_0.3.4, spatstat_1.64-1, rpart_4.1.16, nlme_3.1-157, spatstat.data_2.1-2, rlist_0.4.6.2, scales_1.1.1, forcats_0.5.1, dplyr_1.0.8, tibble_3.1.6, tidyr_1.2.0, devtools_2.3.2, usethis_2.0.1, SeuratObject_4.0.0.

#### Loaded via a namespace (and not attached)

backports_1.2.1, plyr_1.8.6, igraph_1.2.6, lazyeval_0.2.2, splines_4.1.3, BiocParallel_1.24.1, listenv_0.8.0, scattermore_0.7, digest_0.6.29, htmltools_0.5.2, viridis_0.5.1, fansi_1.0.2, memoise_2.0.0, tensor_1.5, cluster_2.1.3, ROCR_1.0-11, remotes_2.2.0, globals_0.14.0, sandwich_3.0-0, spatstat.sparse_2.0-0, prettyunits_1.1.1, colorspace_2.0-3, xfun_0.22, RCurl_1.98-1.3, callr_3.5.1, crayon_1.5.0, jsonlite_1.7.2, zoo_1.8-9, glue_1.6.2, polyclip_1.10-0, gtable_0.3.0, zlibbioc_1.36.0, XVector_0.30.0, leiden_0.3.7, DelayedArray_0.16.2, pkgbuild_1.2.0, future.apply_1.7.0, abind_1.4-5, pheatmap_1.0.12, futile.options_1.0.1, miniUI_0.1.1.1, Rcpp_1.0.6, viridisLite_0.3.0 xtable_1.8-4, reticulate_1.18, spatstat.core_2.0-0, htmlwidgets_1.5.4, httr_1.4.2, ellipsis_0.3.2, ica_1.0-2, pkgconfig_2.0.3, uwot_0.1.10, deldir_0.2-10, utf8_1.2.2, tidyselect_1.1.2, rlang_1.0.2, reshape2_1.4.4, later_1.1.0.1, munsell_0.5.0, tools_4.1.3, cachem_1.0.4, cli_3.2.0, generics_0.1.2, ggridges_0.5.3, evaluate_0.14, stringr_1.4.0, fastmap_1.1.0, yaml_2.3.5, goftest_1.2-2, processx_3.4.5, knitr_1.31, fs_1.5.0, fitdistrplus_1.1-3, RANN_2.6.1, pbapply_1.4-3, future_1.21.0, mime_0.10, formatR_1.8, compiler_4.1.3, plotly_4.9.3, png_0.1-7, testthat_3.0.2, spatstat.utils_2.3-0, stringi_1.5.3, ps_1.6.0, desc_1.3.0, lattice_0.20-45, vctrs_0.3.8, pillar_1.7.0, lifecycle_1.0.1, spatstat.geom_2.0-1, lmtest_0.9-38, RcppAnnoy_0.0.18, bitops_1.0-6, data.table_1.14.2, cowplot_1.1.1, irlba_2.3.3, httpuv_1.5.5, patchwork_1.1.1, R6_2.5.1, promises_1.2.0.1, KernSmooth_2.23-20, gridExtra_2.3, parallelly_1.24.0, sessioninfo_1.1.1, codetools_0.2-18, lambda.r_1.2.4, pkgload_1.2.0, rprojroot_2.0.2, withr_2.4.1, sctransform_0.3.2, GenomeInfoDbData_1.2.4mgcv_1.8-40, rmarkdown_2.7, Rtsne_0.15, shiny_1.6.0.

### Quality control of cells and samples

For each library included in this study, we excluded low quality cells (poor sequenced, damaged or dead cells) by filtering out cells with < 1,000 reads or < 100 genes detected; We also excluded cells with > 15% of reads aligning to mitochondrial genes. Doublets were firstly excluded by removing cells with a number of reads or genes more than three standard deviations from the library mean (Extended Data Fig.2a). Doublets removal was further refined with cell hashing information by using *Seurat*’s function *MULTIseqDemux()* to assign cells to either singlets, doublets or negatives (Extended Data Fig. 2b). Only singlets were retained, as negative cells cannot be demultiplexed and assigned with certainty to the sample of origin. For one library out of 20 sequenced for this study (library 9) the counts related to one hash (sample MG452) were high across all samples in the library and *MULTIseqDemux()* failed to demultiplex many cells. We used the function *HTODemux()* instead to demultiplex library 9 which performed better. Sample MG452 was entirely removed in further QC steps (see below). Genes detected in less than three cells were excluded from the count matrices. At this step, when QC of single cells was completed, the dataset consisted of 154,624 cells across 101 independent mice and 20 sequencing libraries (Extended Data Fig. 2d, c and Table S3). For detailed sequencing statistics per library see Supplementary Tables.

### Normalisation and integration

After QC, each library was individually normalised and scaled using *SCTransform()*. For all the libraries, we selected the 3000 most variable features for downstream integration. We determined a list of common integration anchors across libraries with *FindIntegrationAnchors()* that we used as an input for integration. To integrate all the libraries, we used the *IntegrateData()* function of the *Seurat* package to correct for any potential library batch effect. Integrated matrix was used for downstream analysis (Extended Data Fig. 2c, d).

In the integrated dataset, we performed Principal Components analysis (PCA) and found that the highest variability in the dataset was explained by the separation of CAMs from the microglial cell states, while integrated sequencing libraries did not show any batch effect in the dataset (Extended Data Fig. 2c, d). We selected 27 dimensions for dimensionality reduction by Uniform Manifold Approximation and Projection (UMAP), that we performed with the *RunUMAP()* function. To produce the initial UMAP as in Extended Data Fig. 3a we used the following parameters in *RunUMAP()*: *dims = 1:27, n*.*neighbors = 30L, n*.*epochs = 200, min*.*dist = 0*.*01*. To identify clusters, we used first the function *FindNeighbors()* (parameters for Extended Data Fig. 3a: *dims = 1:20, k*.*param = 100, nn*.*method = “annoy”, annoy*.*metric = “cosine”*) and then performed unbiased clustering by using *FindClusters()* (parameters for Extended Data Fig. 3a: *resolution = 0*.*6, n*.*iter=1000, n*.*start = 10, algorithm = 1, group*.*singletons = T*). This led to the identification of 14 clusters, 10 of which represented unique microglial/myeloid cell states identities, 3 were merged into one homeostatic cluster for their overlapping transcriptomic signature, and 1 resulted in a small low-quality cells cluster. The specific parameters used for UMAP and clustering were defined after assessment of a wide range of possible parameters, which were evaluated in light of cell state annotations and differential expression (DE) results. We start from underclustering and we progressively increase the resolution by identifying further heterogeneity in the data, but we prevent overclustering by assessing that high resolutions lead to the definition of extra clusters that do not significantly differ in gene expression from the existing ones. Cell states were annotated by means of DE (*FindAllMarkers()* function for overall DE, *FindMarkers()* for side by side comparison) and using the *AddModuleScore()* function with a large number of published datasets, GO categories, pathways and signatures as in input^4,7,8,10,15,21^ (Extended Data Fig. 3b). Out of 14 clusters, 9 clustered together and showed high expression of human microglia genes (*P2RY12, CX3CR1, P2RY13, etc*). 3 of these clusters were merged into a Homeostatic Microglia (HM) cell state for their common signature, while 6 unique cell states were assigned to the other microglia clusters (Extended Data Fig. 3b, c, see main text and Fig. 1b, c, d for details). The remaining 5 clusters were enriched in non-microglia markers. Out of those, 1 small population clustered away from the main microglia clusters and expressed high levels of CNS-associated macrophages (CAM) markers (9645 cells, marked by *CD163, MRC1, RNASE1* – Extended Data Fig. 3c, d) as previously described by Mancuso et al.^15^. 3396 cells clustered apart and expressed proliferation markers (high in *TOP2A, MKI67, STMN1* – Extended Data Fig. 3c, d). Although the dataset was previously QCed, a population of Low Quality/Doublets (3517 cells) was still present. A small cluster of secretory cells was defined by sharing the “secretory” signature previously identified also in Hasselman et al.^8^ (1156 cells, high in *AGR2, MNDA* – Extended Data Fig. 3c,d). Finally, we defined a small cluster of other myeloid cells characterised by low expression of microglia homeostatic genes and expression of macrophage/monocyte markers (*CD14, NEAT1, MAFB*) and pro-inflammatory markers (*IL1B, CCL2, CCL3*) (3078 cells – Extended Data Fig. 3c). The top DE genes from each defined cell state confirmed the unique signatures and no over-clustering (top 20 marker genes per cluster are visualised in Extended Data Fig. 3c).

### Microglia subsetting, re-clustering and annotation

For the final analysis of high quality isolated human microglia, we subsetted out other myeloid and low quality clusters (CAM, Other Myeloid, Secretory, Proliferating and Doublets/LowQuality). After trimming, 127,755 human microglia were retained for downstream analysis. Using the previously integrated dataset, we performed PCA and selected 30 dimensions for dimensionality reduction by UMAP, as described above. No library-dependent batch effect was observed (Extended Data Fig. 2d). To produce the final UMAP as in Figure 1b we used the following parameters in *RunUMAP()*: *dims = 1:30, n*.*neighbors = 30L, n*.*epochs = 500, min*.*dist = 0*.*05*. To identify clusters, we used first the function *FindNeighbors()* (parameters for Figure 1b: *dims = 1:20, k*.*param = 100, nn*.*method = “annoy”, annoy*.*metric = “cosine”*) and then performed unbiased clustering by using *FindClusters()* (parameters for Figure 1b: *resolution = 0*.*7, n*.*iter=1000, n*.*start = 10, algorithm = 1, group*.*singletons = T*). This led to the identification of 11 clusters, 4 of which were merged into a Homeostatic Microglia (HM) cell state for their overlapping transcriptomic signature (Figure 1b,c and Extended Data Fig. 3c), and the other 7 represented cell states defined by unique or transitory profiles. With the higher resolution provided by the microglia subclustering, we could identify 2 distinct CRM populations (CRM-1 and CRM-2) that were initially grouped together (Extended Data Fig. 3a,c) but actually represent two consecutive stages on the same phenotypic trajectory, which can be differentially modified by Aβ pathologies and genetic backgrounds (Figure 1b,d, Figure2c,d, Figure 4g,h,I, Figure 5b,c). The specific parameters used for UMAP and clustering were defined after assessment of a wide range of possible parameters, which were evaluated in light of cell state annotations, DE results and samples distribution. Cell states annotation was performed as described for the full dataset, by means of iterative clustering, DE and signature scores (Figure 1c and Extended Data Fig. 4b). We finally defined 8 microglial cell states that included Homeostatic (HM), Cytokine response-1 and 2 (CRM-1 and CRM-2), Interferon response (IRM), Disease Associated (DAM), Antigen-presenting response (HLA), Ribosomal microglia (RM) and Transitioning CRM (Figure 1b, c, d). The expression profiles of top DE genes from each defined cell state (top 10 markers on heatmap in Figure 1d, top 3 markers on UMAP in Extended Data Fig. 4c and all markers DE statistics in Supplementary Table 5) and the signature scores calculated with *AddModuleScore()* (Extended Data Fig. 4a) confirmed the unique transcriptomic profiles of these clusters and that no over-clustering was performed. We excluded from this dataset 6 mice that showed signs of infection, extremely low cell numbers and/or mice with the vast majority of cells mapping to one unique cell state. The final high-quality microglia dataset consisted of 127,755 cells from 95 independent mice and 20 sequencing libraries (Extended Data Fig. 2d, e). For detailed sequencing statistics per library see Supplementary Table 1. For number of sequenced cells per replicates/conditions and all metadata of this study see Supplementary Table 3 and 4.

### Differential expression

Differentially expressed genes were found by applying the *FindAllMarkers()* function for overall DE and *FindMarkers()* for side by side comparisons, both from the *Seurat* R package. All the reported comparisons in the manuscript were performed with the following parameters: *assay = “SCT”, test*.*use = “wilcox”, min*.*pct = 0*.*01, logfc*.*threshold = 0*.*1*. We used the Wilcoxon rank sum test to calculate P-values. We performed DE on the “SCT” assay calculated from *SCTransform()*, since Pearson residuals resulting from regularized negative binomial regression effectively mitigate depth-dependent differences in differential expression, as described by Hafemeister & Satija 2019^44^. We only tested genes that are detected in a minimum fraction of 1% in either of the two populations. We limited testing to genes which show, on average, at least 0.1-fold difference (log-scale) between the two groups of cells. Only genes with their adjusted *P* value <0.05 (*post-hoc*, Bonferroni correction) were considered as significant.

### Pseudotime

Pseudotime analysis was performed in the final human microglia dataset to infer the phenotypic transitions happening between the different microglial cell states. Unsupervised single-cell trajectory analysis was performed with *Monocle3*, an algorithm that allows to learn the sequence of gene expression changes each cell must go through as part of a dynamic biological process. We used *SeuratWrappers* to convert our microglia *Seurat* object into a *Monocle* object with *as*.*cell_data_set()*. We kept the UMAP embeddings previously calculated with *RunUMAP()* in order to estimate the phenotypic transitions between our annotated cell states. We run *cluster_cells()* and *learn_graph()* (parameters used: *close_loop = T, learn_graph_control = list(rann*.*k=100, prune_graph = TRUE, orthogonal_proj_tip = F, minimal_branch_len = 50)*) to learn the trajectory. To infer how resting microglia transition into reactive cell states, we set the roots of the trajectory with *order_cells()* by selecting the 10 most homeostatic cells in our dataset (based on our previously defined HM signature score), in order to avoid limiting the selection of the root to a manually picked single cell and to account for heterogeneity of the HM cluster. Similar trials that set the origin of the trajectory to different HM cells led to comparable results always identifying the main axes of phenotypic transitions described in Figure 2c.

## Data availability

Data generated in this study are available at Gene Expression Omnibus (GEO) database with accession number that will be provided upon publication. An online platform for user-friendly access and visualisation of the data shown in this study is being built at data.bdslab.org/Mancuso2022. A beta version of the webtool is already available, and will be progressively updated with more data and resources. Other datasets included in the manuscript can be found at GEO (Mancuso et al., GSE137444; Sala Frigerio et al., GSE127893; Hasselman et al., GSE133433; Gerrits et al., GSE148822; Sayed et al., GSE183068; Zhou et al., GSE140511; Keren-Shaul et al., GSE98969; Friedman et al., GSE89482). The list of GWAS risk genes for Figure 6a was obtained according to the criteria of nearest protein coding genes to SNPs as identified in Bellenguez et al. 2022^37^, see Supplementary Table 5. The list of GWAS risk genes for Extended Data Fig. 12a,b was obtained as the union of candidate genes from Bellenguez et al., and a candidate gene list selected in our previous publication^22^ (see Supplementary Table 7 for complete list).

## Extended Data Figures

**Extended Data Fig. 1.**
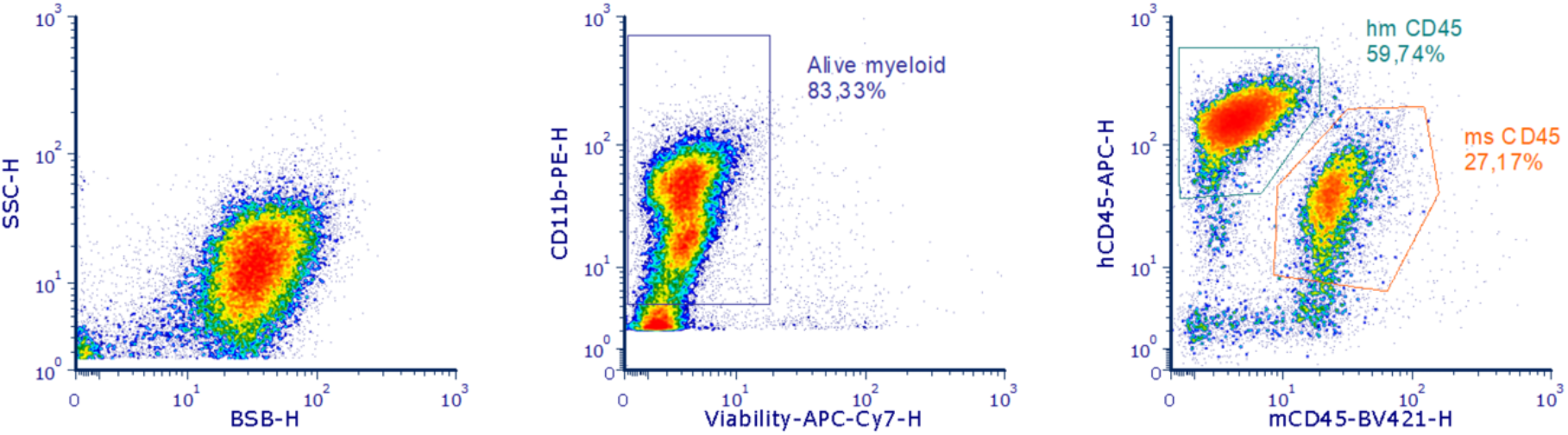
Gating strategy for the isolation of xenotransplanted human microglia from the mouse brain. Human cells were sorted according to the expression of CD11b+ hCD45+ whereas mouse cells were defined as CD11b+ hCD45-mCD45+. Note that we applied a trigger (=threshold) on CD11b+ events, therefore all CD11b negative cells are not displayed.

**Extended Data Fig. 2.**
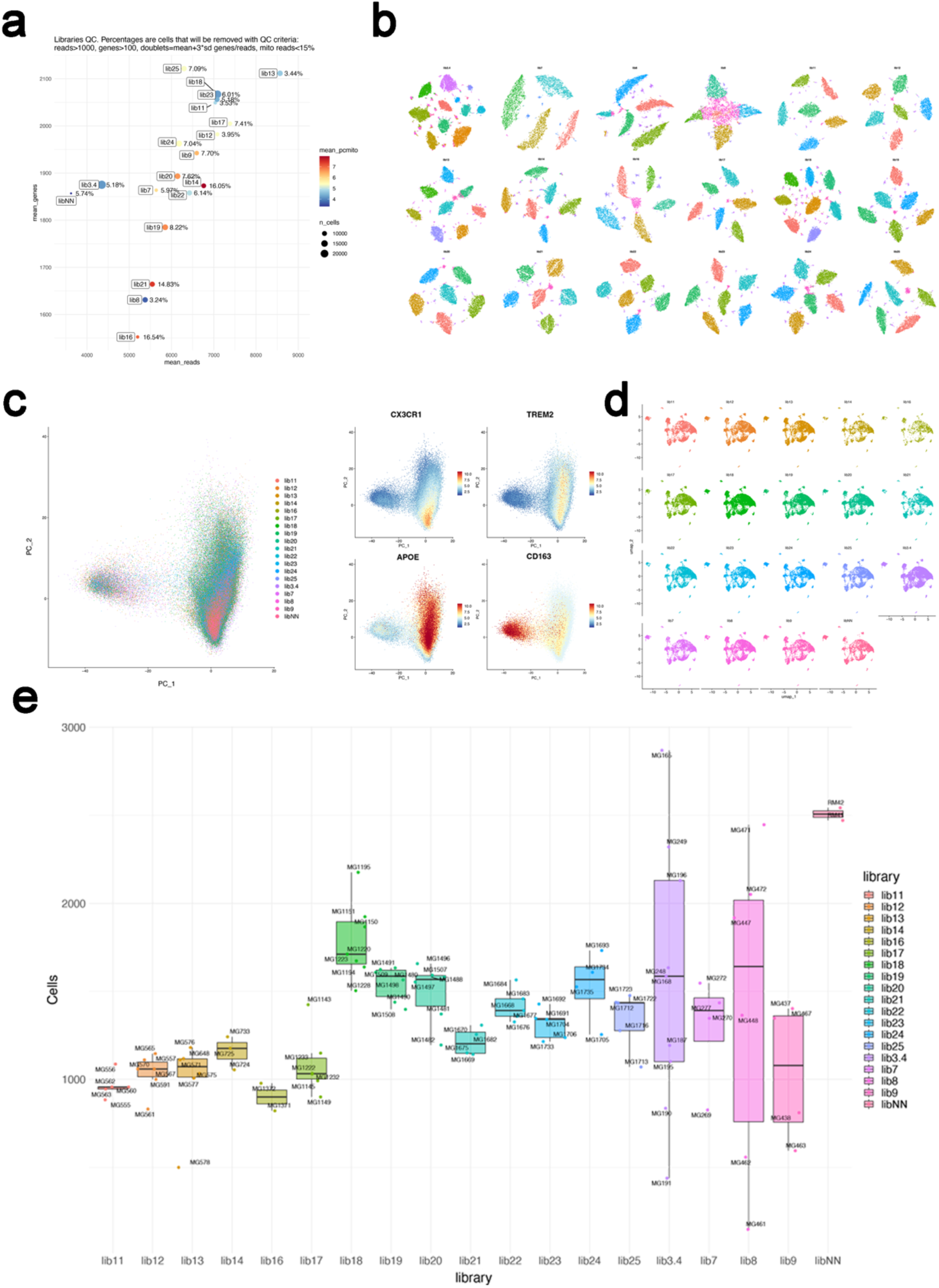
Overview of all the cDNA libraries used in single cell RNA-seq experiments. **a**, Basic quality control of the libraries including number of sequenced cells (size of the dot), mean number of genes (x-axis) and reads (y-axis), as well as mean percentage of reads mapping to mitochondrial genes (colour of the dot). Percentage labels indicate the proportion of cells per library that did not pass QC based on reported criteria (see also Methods). **b**, Cell-hashing counts-based tSNE plots showing the total-seq A barcoded antibody demultiplexing. Each tSNE represents a library, whereas each cluster is an individual mouse assigned to a cell hash. Small clusters are unassigned cells or remaining doublets from previous QC. **c**, First and second principal component (PC) used for the clustering analysis shown in Extended Data Fig. 3a. Libraries completely overlap after data integration, and most of the dataset variability (PC1) is explained by the separation of CAMs (*CD163*) from microglia clusters (*CX3CR1, TREM2, APOE*) (see Extended Data Fig. 3a and Methods). **d**, Individual UMAP plots as in Extended Data Fig. 3a showing the distribution of cells from each single library. **e**, Final number of cells per mouse and per library. Each box represents a library, whereas each dot represents an individual mouse replicate (labelled with MG unique IDs, lib NN corresponds to the data set in Mancuso et al. 2019^15^).

**Extended Data Fig. 3.**
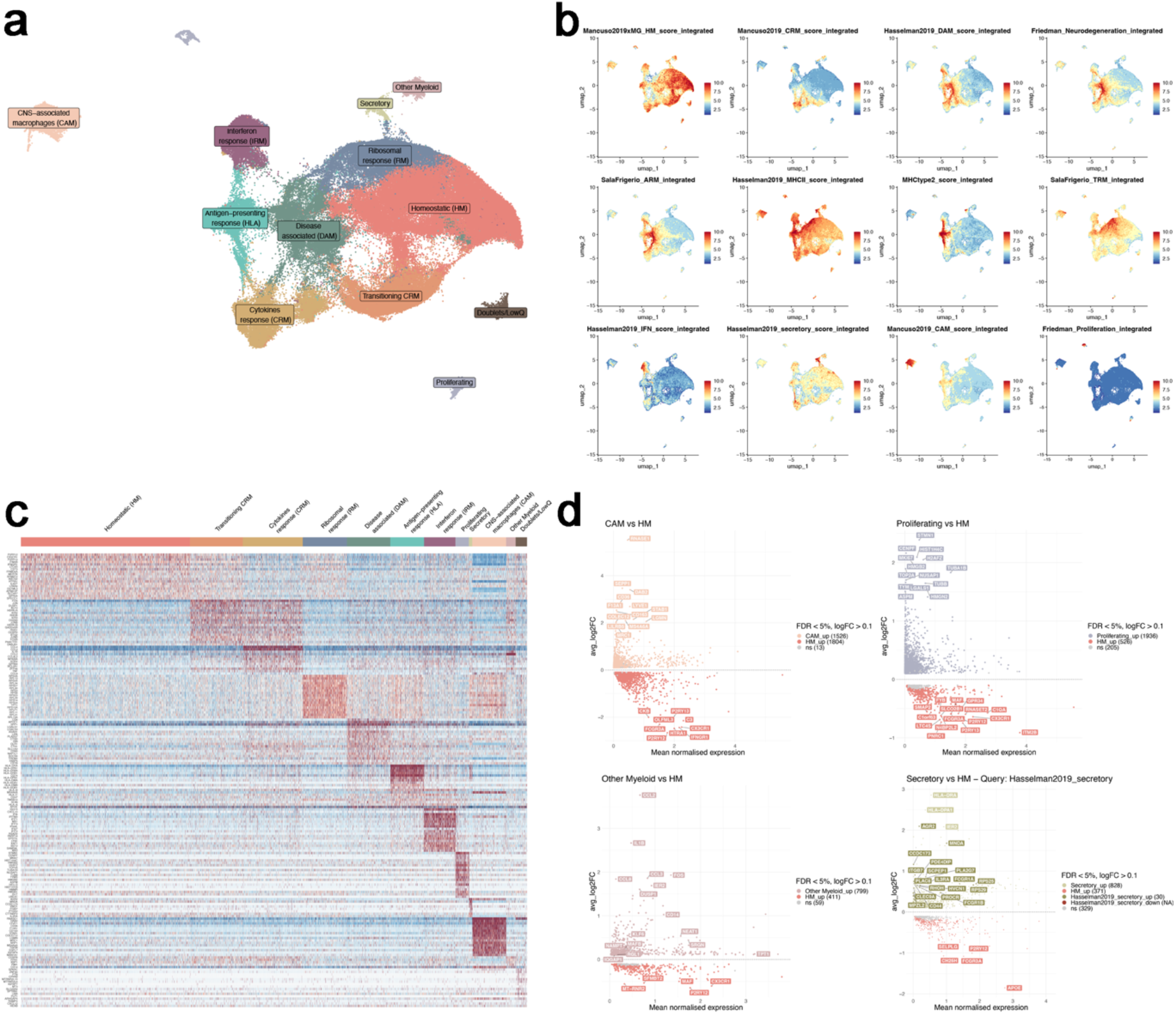
Extended clustering, preparation of the datasets for analysis and cell type/state annotations. **a**, UMAP plot of the 154624 cells passing quality control, coloured by annotated cell states before removal of CAMs, other myeloid, low quality and proliferating clusters. **b**, UMAP plots as in **a**, coloured by the combined level of expression of groups of genes that characterise distinct microglial and peripheral cell states. **c**, Heatmap displaying the top20 most upregulated genes in each cluster as in **a. d**, MA plots with direct comparisons between clusters excluded from the analysis and homeostatic microglia (HM).

**Extended Data Fig. 4.**
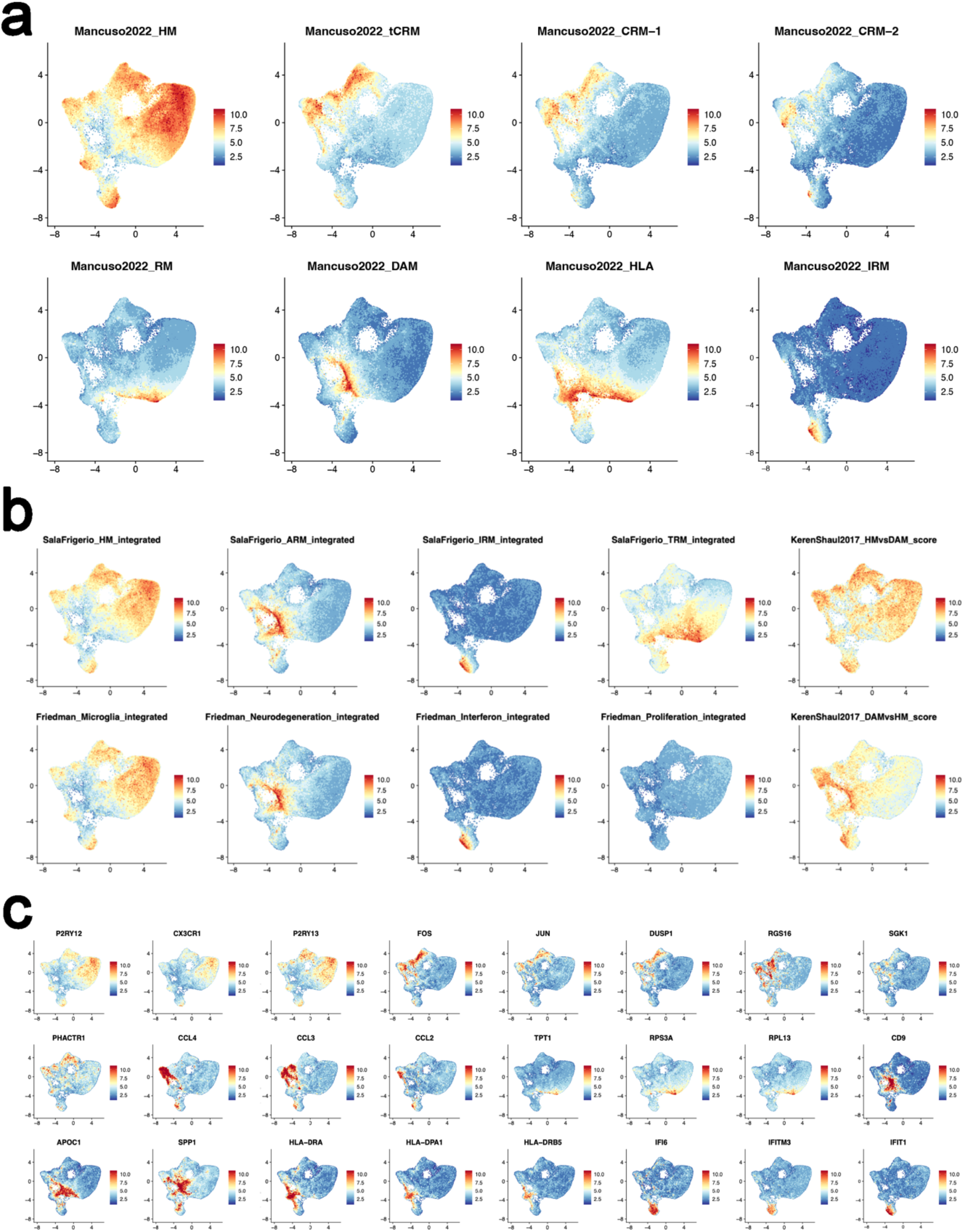
Extended cell state annotations. **a**, UMAP plots as in **Fig. 1a**, coloured by the combined level of expression of groups of genes that characterise distinct microglial states described in this study. **b**, UMAP plots as in Fig. 1a coloured by the combined level of expression of groups of genes that characterise distinct cell states described in mouse studies. **c**, Selected genes defining the different transcriptomic signatures shown in Fig. 1b (3 markers per cell state).

**Extended Data Fig. 5.**
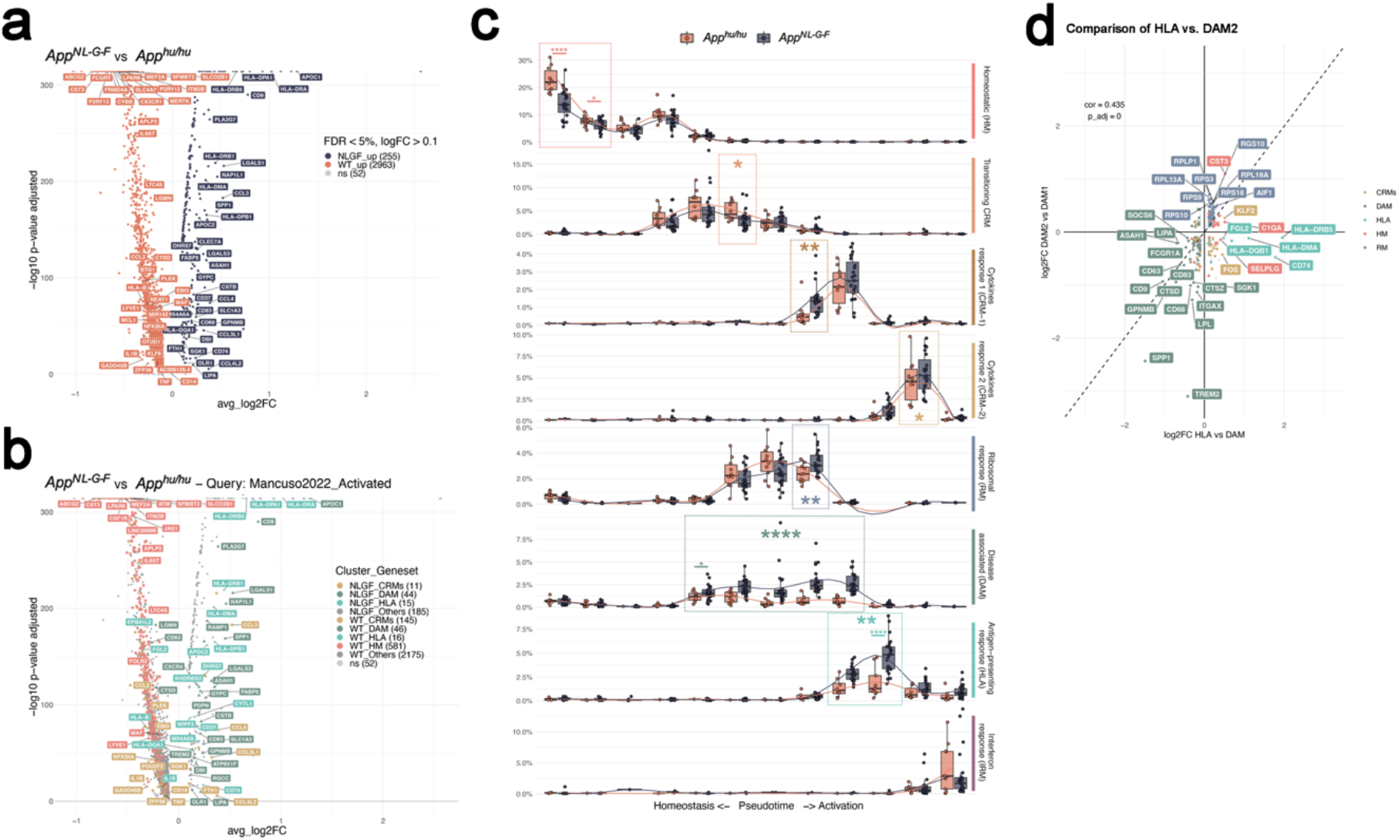
Extended analysis of the cell state transition in *App*^*NL-G-F*^ and *App*^*hu/hu*^ mice. **a**, Volcano plot showing a paired comparison between *App*^*NL-G-F*^ and *App*^*Hu/Hu*^ mice (Wilcoxon rank-sum test, P-values adjusted with Bonferroni correction based on the total number of genes in the dataset). The number of significant genes per condition is reported in brackets in the legend (Log2 Fold-Change threshold > 0.1). Adjusted p-value threshold < 0.05 (ns = not significant). **b**, Volcano plot showing a paired comparison between *App*^*NL-G-F*^ and *App*^*Hu/Hu*^ mice (Wilcoxon rank-sum test, P-values adjusted with Bonferroni correction based on the total number of genes in the dataset), coloured by clusters as in Fig. 1 (HM = red squares, CRMs (CRM-1 & CRM-2) = ochre circles, DAM = olive triangles, HLA = cyan diamonds, other markers = grey circles). The number of significant genes per condition is reported in brackets in the legend (Log2 Fold-Change threshold > 0.1). Adjusted p-value threshold < 0.05 (ns = not significant). **c**, Phenotypic trajectory followed by the human microglia obtained by an unbiased pseudotime ordering with *Monocle 3*. Proportion of cells (y-axis) present in each cell state as in Fig. 1a (rows) at different stages of the pseudotime trajectory (x-axis) in *App*^*NL-G-F*^ and *App*^*hu/hu*^ mice. The data is displayed as box plots, showing all individual mouse replicates for each pseudotime bin. **d**, Correlation analysis of the logFC in DAM2 vs. DAM1 clusters defined by Keren-Shaul et al.^4^ (y-axis) and HLA vs. DAM clusters defined in this study (x-axis). Pearson’s correlation, R= 0.44, differentially expressed genes were adjusted using Bonferroni correction and colored according to clusters in Fig. 1a. Note the specific shift in HLA markers on the x-axis, while DAM2 actually downregulates DAM markers to overexpress mainly ribosomal genes.

**Extended Data Fig. 6.**
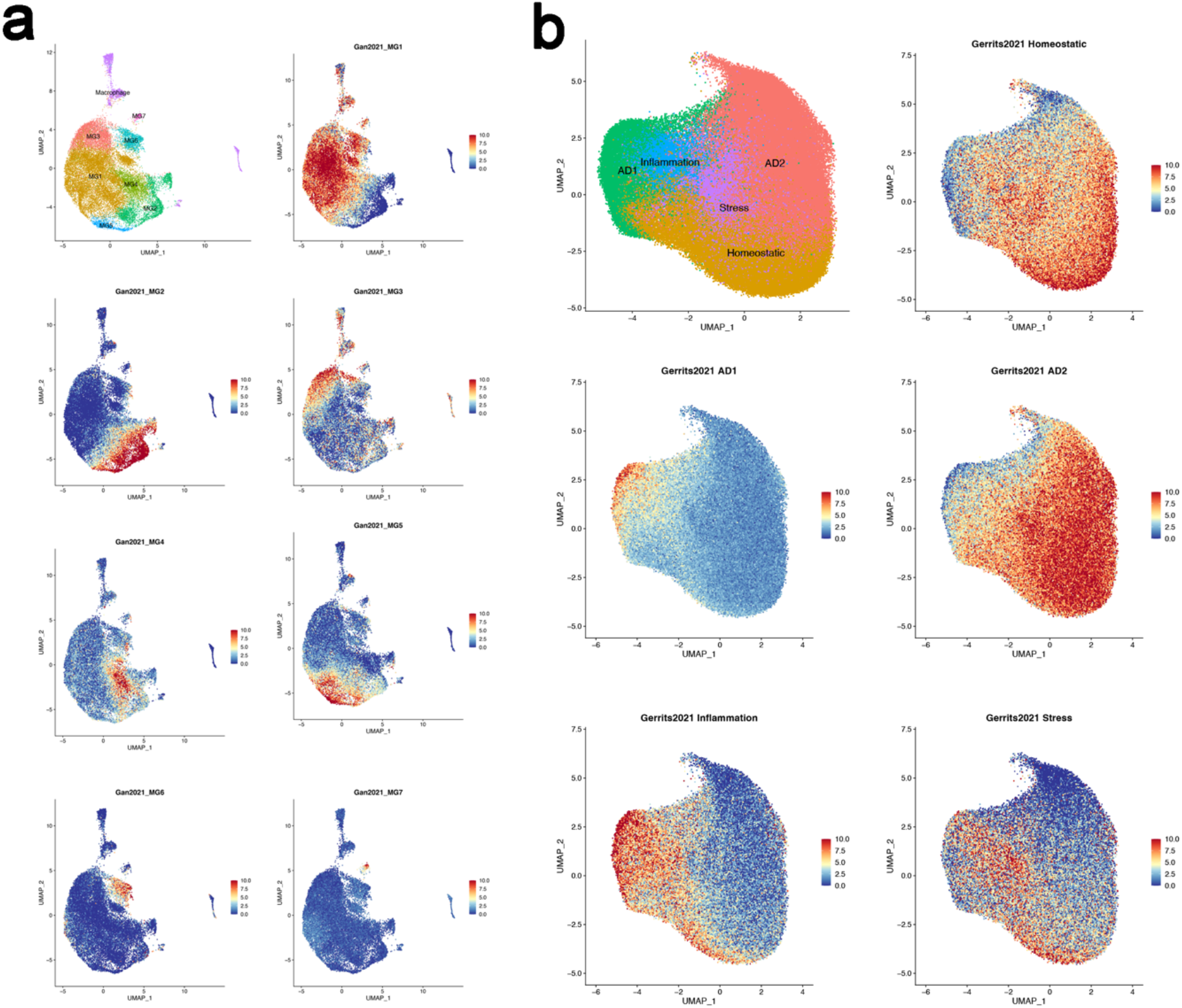
Extended analysis of the single microglia nuclei from human post-mortem brains. **a**, Seurat Module scores of sub-cluster genesets from supplementary data provided by Sayed et al. 2021 ran against re-clustered Sayed et al. 2021 single nuclei data following analysis workflow as described to reproduce annotations from the original study (see Methods, Sayed et al. 2021). **b**, Seurat Module scores of sub-cluster genesets from data provided by Gerrits et al. 2021 ran against re-clustered Gerrits et al. 2021 single nuclei data following analysis workflow as described to reproduce annotations from the original study (see Methods, Gerrits et al. 2021).

**Extended Data Fig. 7.**
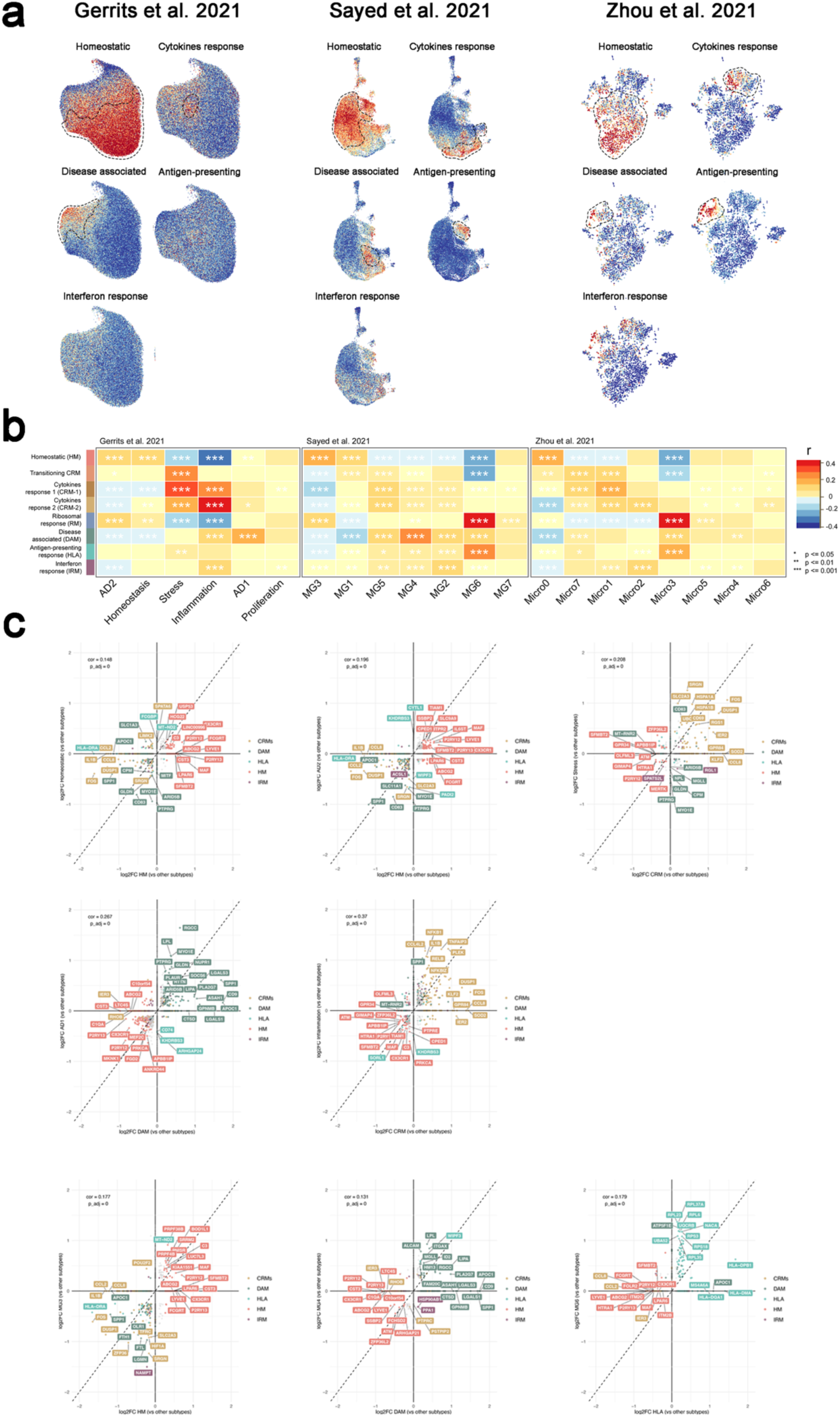
Extended analysis of the single nuclei from human post-mortem brains (continuation). **a**, UMAP plots as in Fig. 5, coloured by the combined level of expression of groups of genes that characterise distinct microglial transcriptional states from xenotransplanted microglia. **b**, Full set of pairwise Pearson correlations between logFC of all DE genes (P-value < 0.05) of each microglia sub-type and logFC of all DE genes (p<0.05) of clusters from each human snRNA-seq study, with significance denoted by asterisks (* p<=0.05, ** p<=0.01, *** p<=0.001). **c**, Representative scatter plots showing the correlation analysis of the logFC between xenotransplanted and single nuclei postmortem microglia from Sayed et al. and Gerrits et al. The labels are coloured by cell state as in Fig. 1.

**Extended Data Fig. 8.**
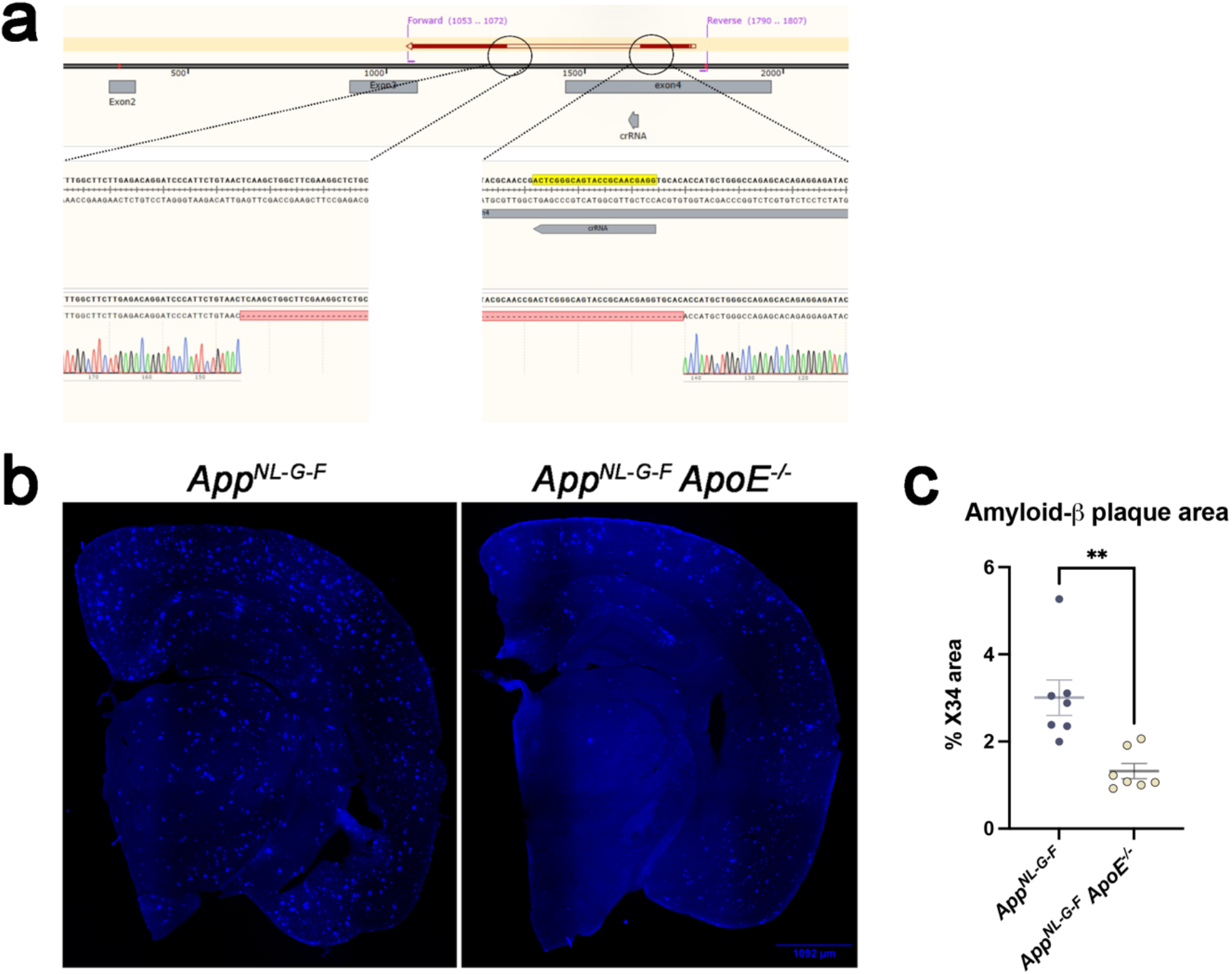
Reduction of amyloid-β plaque in ApoE^-/-^ mice. **a**, Schematic representation of the generation of *App*^*NL-G-F*^ *ApoE*^*-/-*^ mice. The upper part represents the mouse *ApoE* allele on chromosome 7. Exons are indicated by grey boxes. The position of the start codon in exon 2 and the stop codon in exon 4 are indicated in red. The position of the primers used for genotyping are highlighted in pink. The sequence targeted by the Cas9 is labelled in yellow. The lower part shows the Sanger sequencing validation, with a confirmed deletion of 335bp spanning the last 148bp of intron 3 and first 187bp of exon 4. **b**, Representative coronal sections of *App*^*NL-G-F*^ (n=7) and *App*^*NL-G-F*^*xApoE*^*-/-*^ (n=7) mice stained with X34. **c**, Quantification of the % X34 area (unpaired t-test, ** p<0.01)

**Extended Data Fig. 9.**
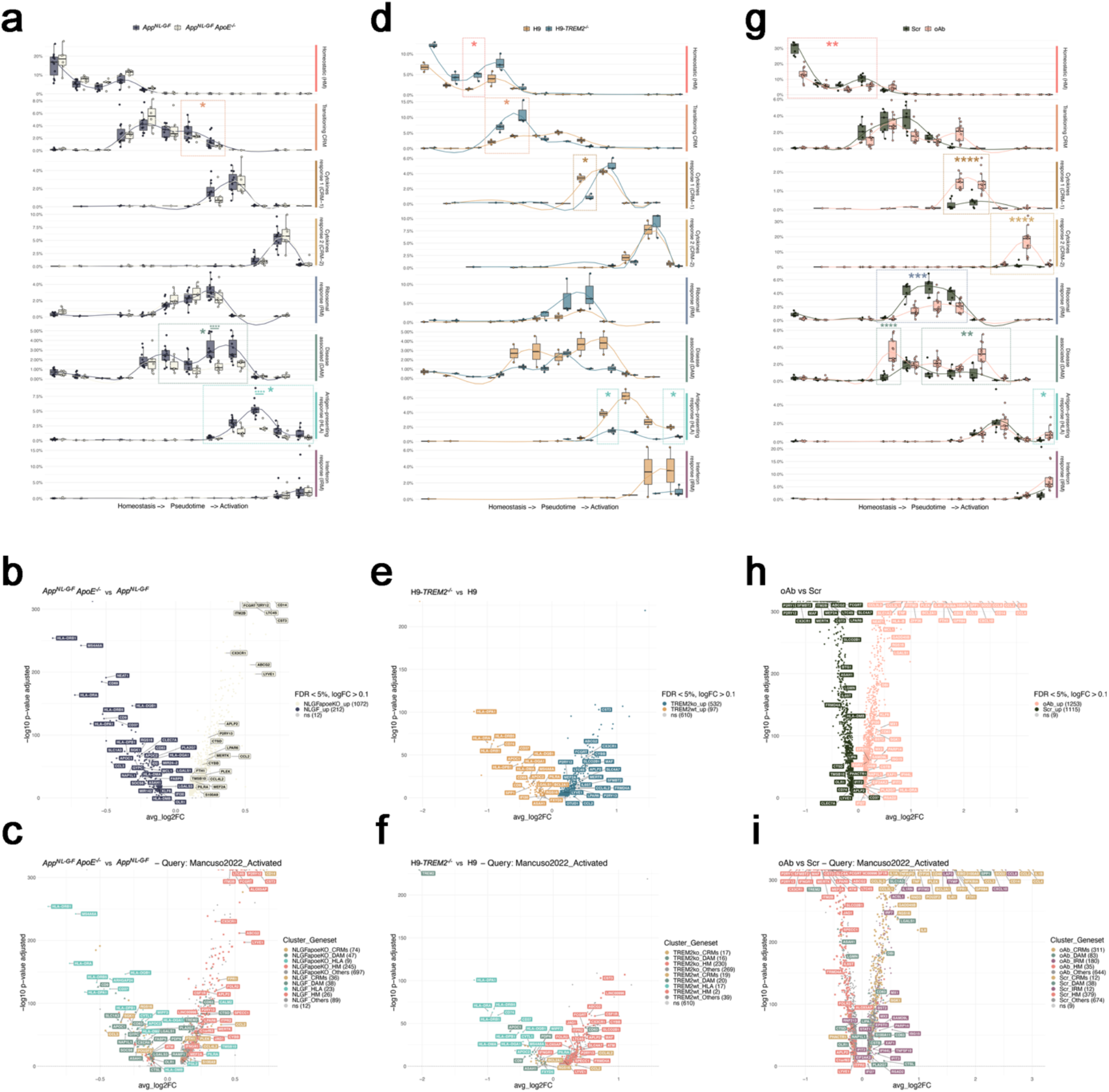
Extended analysis on the differential effects of amyloid-β plaques and soluble aggregates on microglia cell states. **a** Phenotypic trajectory followed by the human microglia obtained by an unbiased pseudotime ordering with *Monocle 3*. Proportion of cells (y-axis) present in each cell state as in Fig. 1a (rows) at different stages of the pseudotime trajectory (x-axis), colored by host genotypes. The data is displayed as box plots, showing all individual mouse replicates for each pseudotime bin. **b, c**, Volcano plots showing a paired comparison between microglia transplanted in *App*^*NL-G-F*^ *ApoE*^*-/-*^ and *App*^*NL-G-F*^ mice, coloured by **(b)** host genotype and **(c)** cluster as in Fig. 1a (HM = red squares, CRMs (CRM-1 & CRM-2) = ochre circles, DAM = olive triangles, HLA = cyan diamonds, other markers = grey circles). The number of significant genes per condition is reported in brackets in the legend (Log2 Fold-Change threshold > 0.1). Adjusted p-value threshold < 0.05 (Wilcoxon rank-sum test, P-values adjusted with Bonferroni correction based on the total number of genes in the dataset, ns = not significant). **d**, Phenotypic trajectory followed by the human microglia obtained by an unbiased pseudotime ordering with *Monocle 3*. Proportion of cells (y-axis) present in each cell state as in Fig. 1a (rows) at different stages of the pseudotime trajectory (x-axis), colored by cells genotypes. The data is displayed as box plots, showing all individual mouse replicates for each pseudotime bin. **e, f**, Volcano plots showing a paired comparison between *TREM2*^*-/-*^ and *TREM2*-WT microglia transplanted in *App*^*NL-G-F*^ mice, coloured by **(e)** cells genotype and **(f)** cluster as in Fig. 1a (HM = red squares, CRMs (CRM-1 & CRM-2) = ochre circles, DAM = olive triangles, HLA = cyan diamonds, other markers = grey circles). The number of significant genes per condition is reported in brackets in the legend (Log2 Fold-Change threshold > 0.1). Adjusted p-value threshold < 0.05 (Wilcoxon rank-sum test, P-values adjusted with Bonferroni correction based on the total number of genes in the dataset, ns = not significant). **g**, Phenotypic trajectory followed by the human microglia obtained by an unbiased pseudotime ordering with *Monocle 3*. Proportion of cells (y-axis) present in each cell state as in Fig. 1a (rows) at different stages of the pseudotime trajectory (x-axis), colored by treatment. The data is displayed as box plots, showing all individual replicates for each pseudotime bin. **h, i**, Volcano plots showing a paired comparison between microglia transplanted in *App*^*hu/hu*^ mice and treated with soluble amyloid-β aggregates or scrambled peptide, coloured by **(h)** treatment and **(i)** cluster as in Fig. 1a (HM = red squares, CRMs (CRM-1 & CRM-2) = ochre circles, DAM = olive triangles, IRM = plum diamonds, other markers = grey circles). The number of significant genes per condition is reported in brackets in the legend (Log2 Fold-Change threshold > 0.1). Adjusted p-value threshold < 0.05 (Wilcoxon rank-sum test, P-values adjusted with Bonferroni correction based on the total number of genes in the dataset, ns = not significant).

**Extended Data Fig. 10.**
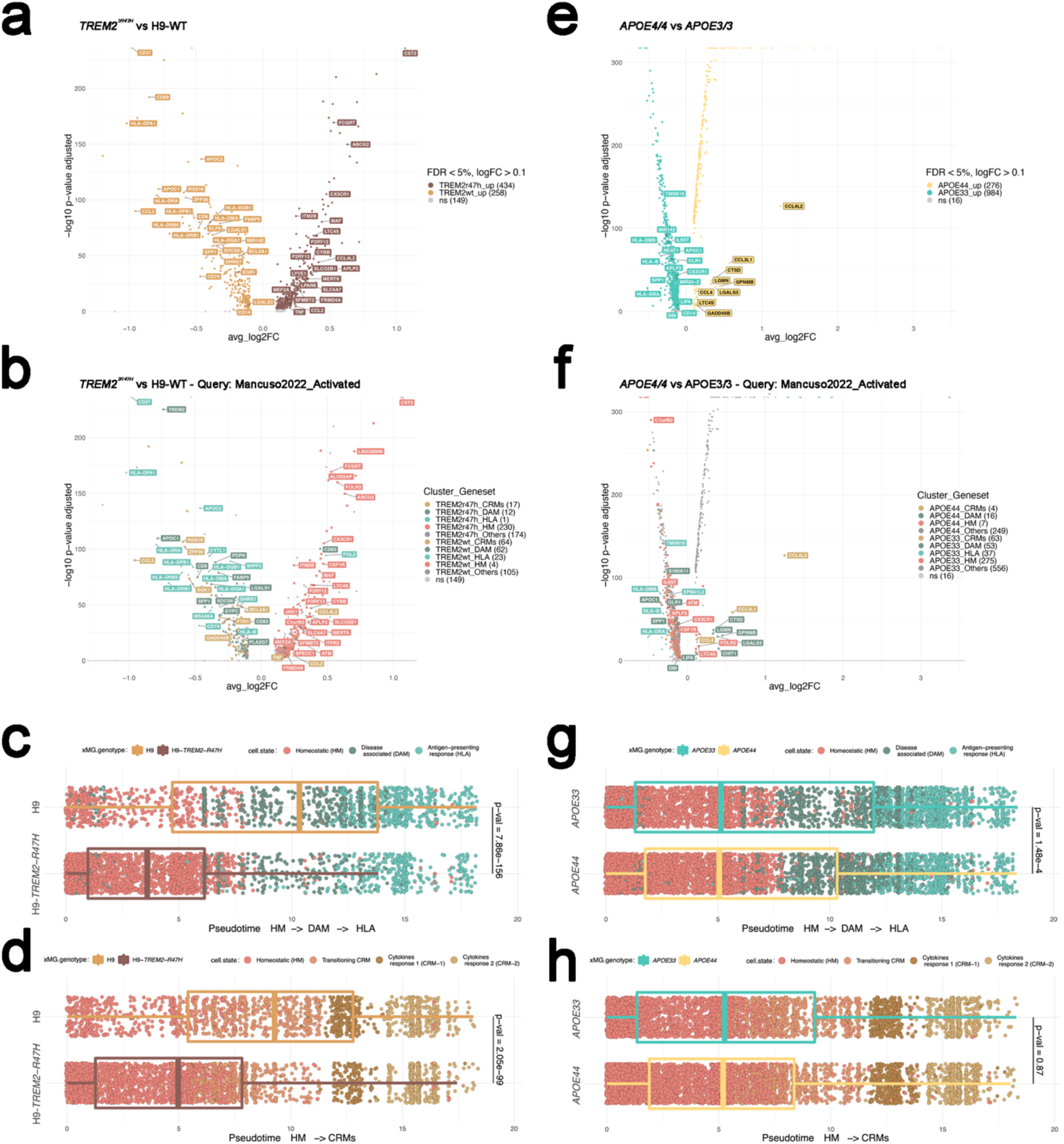
Extended analysis of the impact of clinical mutations on microglial cell states. **a, b**, Volcano plots showing a paired comparison between *TREM2*^*R47H*^ and *TREM2-WT* microglia transplanted in *App*^*NL-G-F*^ mice and coloured by **(a)** cells genotype or **(b)** cluster as in Fig. 1a (HM = red squares, CRMs (CRM-1 & CRM-2) = ochre circles, DAM = olive triangles, HLA = cyan diamonds, other markers = grey circles). The number of significant genes per condition is reported in brackets in the legend (Log2 Fold-Change threshold > 0.1). Adjusted p-value threshold < 0.05 (Wilcoxon rank-sum test, P-values adjusted with Bonferroni correction based on the total number of genes in the dataset, ns = not significant). **c, d**, Distribution of *TREM2*^*R47H*^ and *TREM2-WT* cells across **(c)** DAM-HLA and **(d)** CRM transcriptional trajectories. Note the shift in transcriptional states in both the DAM-HLA and CRM axes. **e, f**, Volcano plots showing a paired comparison between *APOE4/4* and *APOE3/3* microglia transplanted in *App*^*NL-G-F*^ mice and coloured by **(e)** cells genotype or **(f)** cluster as in Fig. 1a (HM = red squares, CRMs (CRM-1 & CRM-2) = ochre circles, DAM = olive triangles, HLA = cyan diamonds, other markers = grey circles). The number of significant genes per condition is reported in brackets in the legend (Log2 Fold-Change threshold > 0.1). Adjusted p-value threshold < 0.05 (Wilcoxon rank-sum test, P-values adjusted with Bonferroni correction based on the total number of genes in the dataset, ns = not significant). **g, h**, Distribution of *APOE4* and *APOE3* cells across **(g)** DAM and HLA and **(h)** CRM transcriptional trajectories. Note the shift in transcriptional states in the HLA axis.

**Extended Data Fig. 11.**
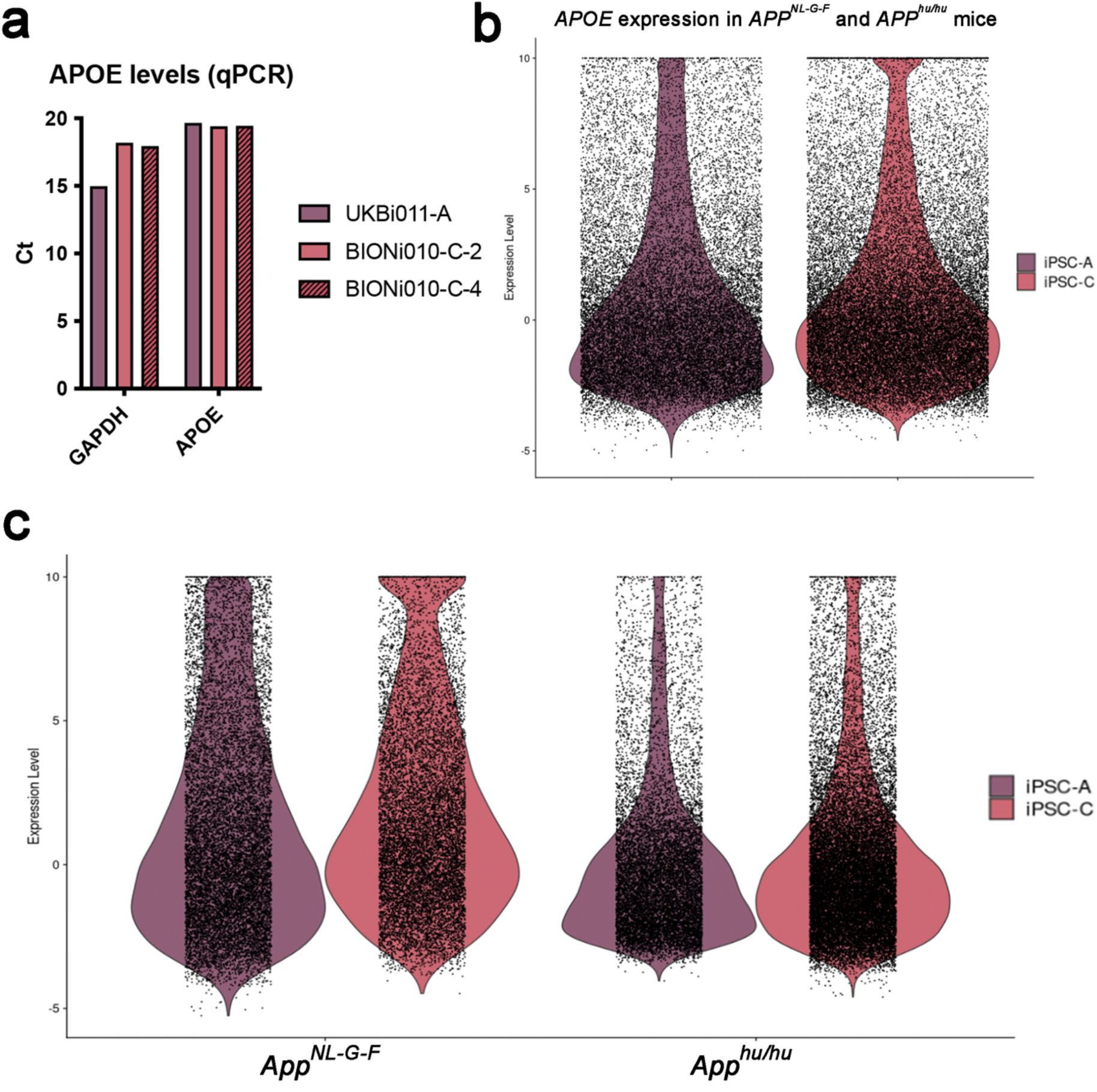
Levels of APOE expression across the different lines used in this study. **a**, Levels of expression of *APOE* and GAPDH in stem cells from UKBi011-A (iPSC-A, e4/e4), BIONi010-C-2 (iPSC-C, e3/KO) and BIONi010-C-4 (iPSC-3, e4/KO). **b**, Aggregated levels of APOE expression across all transplanted microglia used in this study. **c**, APOE expression in human transplanted microglia divided by mouse genotype.

**Extended Data Fig. 12.**
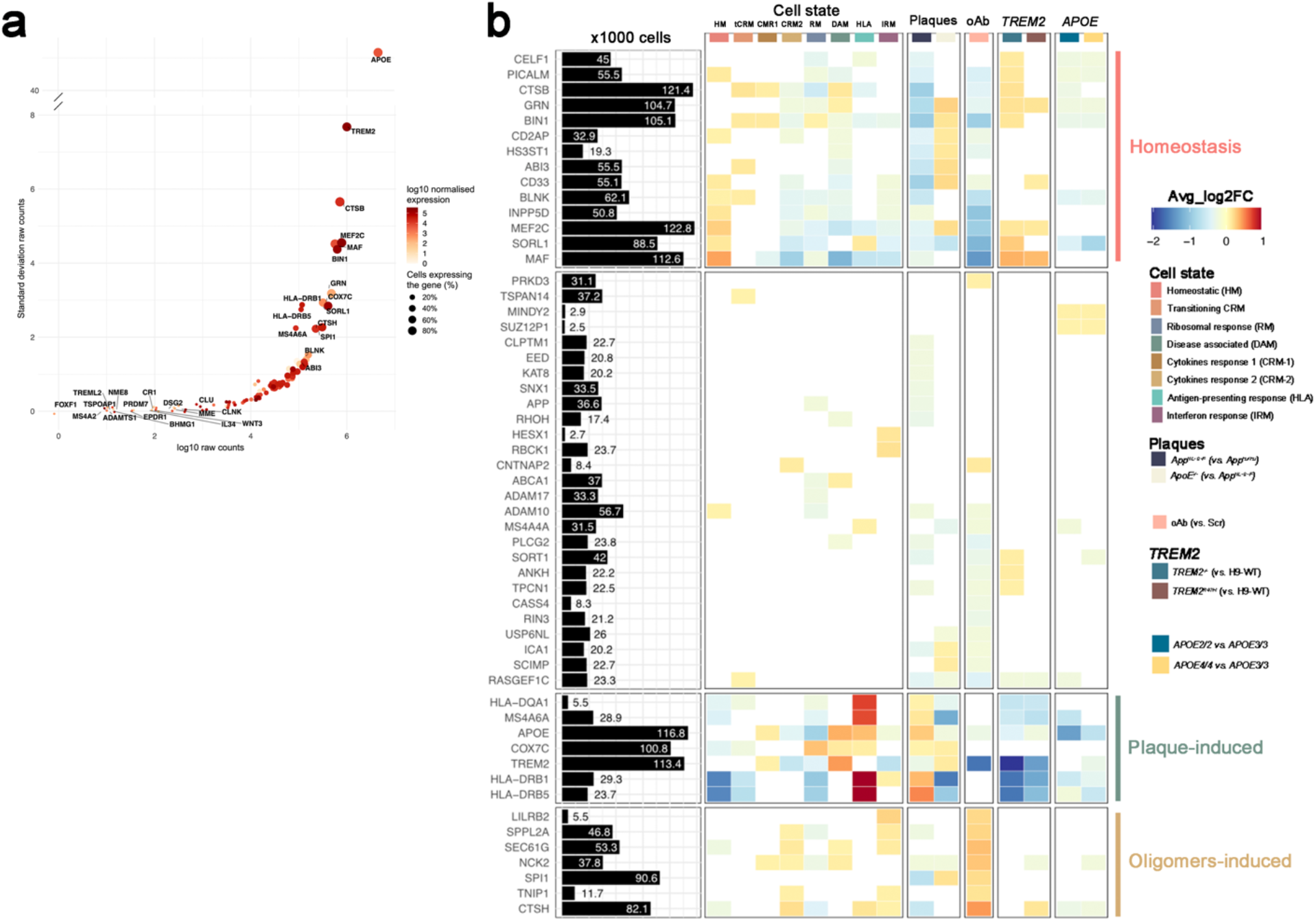
Extended analysis of GWAS genes enrichment in xenotransplanted microglia. **a**, We expanded the gene list with the canonical AD genes according to the latest GWAS study of Fig 6a, with other genes of interest, i.e. the genes selected in our previous publication^22^ (see Supplementary Table 7 for the complete list). Raw counts (log scale) are reported on the x axis, while standard deviation of counts is shown on the y-axis. Dots are coloured by SCT-normalised expression (see Methods) and size reflects percentage of cells in our dataset where the gene is detected. Note that the vast majority of risk genes are detected in our dataset (see all values in Supplementary Table 7). **B** All genes from (a) that are significantly changing in at least one of the tested conditions in this manuscript are reported in an extended heat map build as in Fig.6a. The black bars represent the number of cells (in thousands) with detectable expression (>=1 read per cell) for each candidate gene. The heat map summarizes the deregulated expression (LogFC, colour scale) of these genes across cell states (each cluster compared to all others), as well as after exposure to amyloid-β plaque pathology, upon injection of soluble oligomeric Aβ, or altering the genetic background of the mice or the transplanted cells. The genes are ranked in rows based on hierarchical clustering. We identify 3 sets of genes that display a common profile across cell states (based on their enrichment in the specific microglial phenotypic transcriptional states HM, DAM and HLA, and CRM), amyloid-β pathology and genetic risk, and we group these profiles as: microglia Homeostasis, plaque-induced genes, and oligomers-induced genes. The remaining genes did not show a clear enrichment in cell states or other conditions. All differential expressions were significant after adjusting *P-values* using Bonferroni correction (FDR < 0.05).

